# Intraspecies diversity reveals a subset of highly variable plant immune receptors and predicts their binding sites

**DOI:** 10.1101/2020.07.10.190785

**Authors:** Daniil M Prigozhin, Ksenia V Krasileva

## Abstract

Evolution of recognition specificities by the immune system depends on the generation of receptor diversity, and connecting binding of new antigens with initiation of downstream signalling. In plant immunity, these functions are enabled by the family of innate Nucleotide-Binding Leucine Rich Repeat (NLR) receptors. In this paper we surveyed the NLR complements of 62 ecotypes of *Arabidopsis thaliana* and 54 lines of *Brachypodium distachyon* and identified a limited number of NLR subfamilies responsible for generation of new receptor specificities. We show that the predicted specificity-determining residues cluster on the surfaces of Leucine Rich Repeat domains, but the location of the clusters varies between NLR subfamilies. By comparing NLR phylogeny, allelic diversity, and known functions of the Arabidopsis NLRs, we formulate a hypothesis for emergence of direct and indirect pathogen sensing receptors, and of the autoimmune NLRs. These findings reveal the recurring patterns of evolution of innate immunity and inform NLR engineering efforts.

## Introduction

Plants lack the adaptive immunity of vertebrates. With their immune receptor specificities encoded in the germline, plants can achieve remarkable receptor diversity at the population level (Bakker et al., 2006). The mechanisms that generate this diversity and select for useful and against deleterious receptor variants are thus of great importance to both basic science and to crop improvement (Dangl et al., 2013). Ongoing efforts at pan-genome sequencing of both model and crop species reveal the intraspecies diversity of plant immune receptors, their natural history, mechanisms of action, and the evolutionary forces that shape plant immunity (Van de Weyer et al., 2019; Stam et al., 2019a, 2019b; Seong et al., 2020; Gordon et al., 2017).

Two types of plant immune receptors form the basis of pathogen recognition: extracellular receptors, including receptor-like kinases (RLK) and receptor-like proteins (RLP), and intracellular Nucleotide-binding Leucine Rich Repeat (NLR) proteins (Dangl et al., 2013). While the RLKs and RLPs monitor plant extracellular environments, NLRs are cell death executing receptors shared across plant and animal kingdoms (Jones et al., 2016). Plant NLRs are typically composed of three domains: a central Nucleotide Binding (NB-ARC) domain that mediates receptor oligomerization upon activation, the C-terminal Leucine Rich Repeat (LRR) domain that defines receptor specificity, and one of three N-terminal domains: Resistance To Powdery Mildew 8 (RPW8), Coiled-Coil (CC), or Toll/Interleukin-1 Receptor homology (TIR) domain, these mediate the immune effector function. Based on the N-terminal domains and their evolutionary origin, NLRs are divided into three monophyletic classes: RNL, CNL, and TNL, respectively (Shao et al., 2016).

Functionally, NLRs can be sensors or signal transducers (helpers) (Wu et al., 2017). For example, all RNL genes are thought to be helpers (Jubic et al., 2019), while TNLs and CNLs can fulfill either function. Sensor NLRs have been shown to recognize pathogens using three main modes: i) direct binding to the pathogen-derived effector molecules, ii) indirect recognition of effector activities on other plant proteins, and iii) recognition of modifications to a non-canonical integrated domain of the NLR, which acts as a bait for the effector (Cesari, 2018). The recognition mode of a given sensor NLR is likely to have a large effect on the evolutionary pressure it experiences. Indirect recognition NLRs are likely to undergo balancing or purifying selection based on monitoring a conserved effector activity. In contrast, effector recognition upon direct binding likely requires NLRs to adapt rapidly to keep track of easy-to-mutate effector surface residues. Among the best studied NLRs that directly bind pathogen-derived effectors are the flax L genes (Ellis et al., 2007; Catanzariti et al., 2010), MLA/Sr50 locus in barley and wheat (Chen et al., 2017; Saur et al., 2019), and the RPP1 genes in Arabidopsis (Krasileva et al., 2010; Goritschnig et al., 2016). Their effector targets are structurally diverse, suggesting that individual alleles’ current recognition specificities are recently derived, rather than ancestral.

Continuous generation of diversity in sensor NLRs is required to provide protection from diverse pathogens, and is thought to result from divergent (diversifying) selection and a birth-and-death process acting on NLR gene clusters (Michelmore and Meyers, 1998). NLRs diversify through copy number variation, recombination, gene conversion, gene fusion, and point mutations (Baggs et al., 2017). In a subset of NLRs these mechanisms combine to produce an astounding array of alleles (Bakker et al., 2006; Ding et al., 2007). Not unexpectedly, such diversity comes at a price. Hybrid necrosis has been observed widely in inbreeding and outcrossing plants, cultivated and wild ones and can be considered a plant version of autoimmunity (Bomblies, 2009). Hybrid necrosis occurs due to a mismatch between NLR variants and other plant genes leading to autoimmune recognition as exemplified by Dangerous Mix (DM) genes in *A. thaliana* (Bomblies et al., 2007; Chae et al., 2014; Atanasov et al., 2018) and *Ne* genes in wheat (Zhang et al., 2016). Tomato *Cf-2* is a known example of a non-NLR immune receptor that shows a similar phenotype (Kruger, 2002; Santangelo et al., 2003). These negative interactions revealed in crosses are likely only a small fraction of the cost of derivation of new immune specificities in the presence of the whole intracellular plant proteome.

Cross-species phylogenetic analyses of the NLR gene family have led to important insights into NLR evolution. A combined phylogeny of maize, sorghum, brachypodium, and rice NLRs has been used to identify recently derived NLR immune specificities against rice blast disease (Yang et al., 2013). An expansion of a network of helper and sensor NLRs has been identified across asterids in which a set of diverse sensors signal through a redundant set of helpers that show reduced diversity (Wu et al., 2017). Phylogenetic analyses in grasses have identified major integration clades of NLRs that incorporate additional domains that serve as baits for pathogens (Bailey et al., 2018). However, despite the recent progress in the elucidation of the intra-species NLR complements of both model and non-model plants (Van de Weyer et al., 2019; Stam et al., 2019a, 2019b; Seong et al., 2020; Gordon et al., 2017), systematic analysis of relationships between NLR phylogeny, mode of recognition, and the amount of allelic diversity has not yet been performed.

Recent elucidation of both the pre-activation monomeric and activated resistosome-forming conformations of ZAR1, an indirect recognition CNL, dramatically improved our understanding of both target binding and receptor activation mechanisms of NLRs (Wang et al., 2019b, 2019a). While more NLR structures are likely to be revealed in the future, structure determination efforts will likely lag behind the pan-genome sequencing due to the cost and difficulty of the experiments involved. Therefore, prediction of NLR mode of recognition and of NLR specificity-determining residues from sequence data is an attractive direction that is yet to be fully explored. The idea that highly variable residues determine immune receptor specificity predated the first antibody structure by three years (Kabat, 1970). In the subsequent decades several measures of amino-acid diversity were advanced. Shannon entropy, which originated in information theory, is given by the formula:

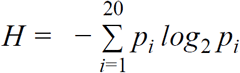

where *p*_*i*_ is the fraction of one of the twenty amino acids in a column of a protein sequence alignment. It was first applied to study antibody and T-cell receptor specificity determining residues (Shenkin et al., 1991; Stewart et al., 1997). High entropy values correlate strongly with surface exposure and hydrophilic character (Liao et al., 2005) and can be used to predict rapidly evolving ligand-binding sites (Magliery and Regan, 2005). In addition to B- and T-cell receptors, entropy-based measures have been applied to identify binding sites in TRP repeat proteins, ankyrin repeat proteins, Zn-finger transcription factors, and G protein-coupled receptors (Sanders et al., 2011; Magliery and Regan, 2005). In this paper, we used phylogenetic analyses to group Arabidopsis and Brachypodium NLRs into near allelic series and applied Shannon entropy analyses of protein alignments to define highly variable NLRs and their candidate specificity determining residues in each species.

Our results show that, depending on the ecotype, 15 to 35 Arabidopsis NLRs belong to rapidly diversifying families. These are distributed in the NLR phylogeny among both CC- and TIR-containing NLRs and encompass the known Dangerous Mix NLRs. We further show that in the highly variable NLRs (hvNLRs), the highly variable residues identified by Shannon entropy cluster on the surface of the LRR domain and contain surface exposed hydrophobic residues surrounded by charged residues, thus, identifying likely binding sites. The exact location of the putative binding sites on the LRR surface is not conserved across different NLRs. Based on the phylogenetic distribution of hvNLRs, we formulate a hypothesis regarding the origin of indirect recognition sensor NLRs. When applied to *Brachypodium distachyon* pan-genome our methods reveal a similarly dispersed phylogenetic distribution of highly variable NLRs in this model grass species. Collectively, our results reveal the origins of novel recognition specificities in NLR innate immune receptors and the common patterns in the evolution of innate immunity.

## Materials and Methods

### Data sources

Arabidopsis pan-NLRome nucleotide assemblies were downloaded from the 2Blades foundation (http://2blades.org/resources/). Gene annotations were downloaded from GitHub pan-NLRome repository (https://github.com/weigelworld/pan-nlrome/). The gene models that matched assemblies were available for 62 *A. thaliana* accessions (Van de Weyer et al., 2019), and these were processed to extract the amino acid sequences of captured protein-coding genes using bedtools getfasta program (Quinlan, 2014). The reference set of 168 NLR alleles (including splice variants) of the Arabidopsis Col-0 genome was extracted as described before (Sarris et al., 2016). Brachypodium proteomes for 54 lines were downloaded from BrachyPan (https://brachypan.jgi.doe.gov) (Gordon et al., 2017).

### Pan NLRome phylogenetic analysis and initial clade assignments

Phylogenetic tree construction for the *A. thaliana* and *B. distachyon* NLRomes and the NLRomes of reference accessions was performed as previously described (Bailey et al., 2018). Briefly, amino acid sequences were searched for the presence of NB-ARC domain using hmmsearch (Mistry et al., 2013) and an extended NB-ARC Hidden Markov Model (HMM) 13059_2018_1392_MOESM16_ESM.hmm (Bailey et al., 2018), and initial alignment was made on this HMM using -A option. The resulting alignment was processed with Easel tools (https://github.com/EddyRivasLab/easel) to remove insertions and retain aligned sequences that matched at least 70% of the HMM model. This alignment was used to construct maximum likelihood phylogeny using RAxML software version 8.2.12 (Kozlov et al., 2019) (raxml -T 8 -n Raxml.out -f a -x 12345 -p 12345 -# 100 -m PROTCATJTT). The trees were visualized in iTOL (Letunic and Bork, 2019). The phylogeny was used to separate protein sequences into clades using R scripts *prefix*_Initial_Assignment.R (hereafter *prefix* is either Atha_NLRome or Brachy_NLRome for the two species under analysis). This and other scripts referenced below are available at (https://github.com/krasileva-group/hvNLR). First, for each NB-ARC sequence a clade of size 40 to 500 with the strongest bootstrap support was chosen. For sequences that did not belong to clades in this size range, smaller clades were allowed. Second, the resulting set of clades was made non-redundant by excluding all nesting clades. The resulting partitions uniquely assigned the 7,818 *A. thaliana* NLR sequences to 65 clades and 11,488 *B. distachyon* NLR sequences to 91 clades.

### Iterative clade refinement

For each identified clade, full length protein sequences were aligned using PRANK algorithm (Löytynoja, 2014) and phylogenetic trees based on full length alignments were constructed as described above using RAxML (Kozlov et al., 2019). Trees were visualized in iTOL, along with subclade statistics calculated in R, and R scripts were used to produce subclade lists based on the trimmed branches (*prefix_*Refinement.R). For the first iteration, gappy columns in the full-length alignments were masked (90% cutoff), later iterations were analyzed without masking gappy columns. Clade refinement was performed as follows: all tree branches longer than 0.3 were cut to form 2 or more subclades. All branches 0.1 and shorter were retained in the first iteration, and for the branches between 0.1 and 0.3, the decision to cut was made by visually inspecting the tree in iTOL and considering bootstrap support and overlap in ecotypes on either side of a branch. The sequences belonging to the refined subclades were realigned using PRANK and tree construction repeated. In the following iterations some branches shorter than 0.1 were cut based on tree inspection in iTOL based on bootstrap support and ecotype overlap. All clade trees generated during this project are available at http://itol.embl.de/shared/daniilprigozhin. The refinement process converged to produce the final assignment of all genes into 237 final clades for *A. thaliana* and into 433 clades for *B. distachyon*.

### Identification of hvNLR clades and binding site prediction in hvNLRs

We used R scripts (*prefix_*CladeAnalysis.R) to calculate alignment Shannon entropy scores using package “entropy”. Alignments that contained 10 or more positions with at least 1.5 bits were considered highly variable. All highly variable clades were examined for the presence of Arabidopsis Col-0 allele. For these Col-0 alleles, we predicted the LRR coordinates manually and cross-checked these predictions with a LRRpredictor online server (Martin et al., 2020). R script was used to map entropy scores to the predicted concave surface of the LRR domain (Atha_NLRome*_*GeneEntropy.R). The entropy scores for the individual strands of LRRs (LxxLxLxx) were exported in tabular format. The hydrophobicity scores for these residues were calculated as the percent of hydrophobic residues at a given amino acid position and exported as a second table. Resulting 2D representations of entropy and hydrophobicity of the concave sides were visually examined for clustering of residues that showed both high entropy scores and the presence of hydrophobic residues. Positive selection analysis of RPP13 clade alignment was carried out in PAML (Yang, 2007).

### Comparison of RPP13 homology model and ZAR1 structure

In order to compare the 3D spatial distribution of highly variable residues in RPP13 with the ZAR1-RKS1 binding site, we used phyre2 in one-to-one threading mode to produce a model for RPP13 (Kelley et al., 2015). Crucially for the accuracy of the model, there are few gaps in the alignment and these gaps are small. R script (Atha_NLRome_GeneEntropy.R) was used to produce a Chimera-formatted attribute file to color the model surface by entropy scores and figure was generated in Chimera (Pettersen et al., 2004).

## Results

### Arabidopsis NLRome shows variable rates of NLR diversification

Recent elucidation of the NLR complements of over 60 accessions of the model plant *A. thaliana* (Van de Weyer et al., 2019) provided a unique opportunity to examine rapidly evolving clades of Arabidopsis NLRs. Unique advantage of the Arabidopsis dataset is the ability to correlate observed diversity to known functional classes of the extensively characterized NLRs. Previous NLRome analyses on this dataset were performed using OrthoMCL followed by orthogroup refinement, and while providing a valuable base for global analyses of selection pressures, did not produce robust allelic series for each gene. This was likely caused by the divergent rates of diversification across NLRs, which complicate orthogroup assignment. To circumvent this challenge, we adopted a phylogeny-based approach. To group NLRs into near allelic series, we first built a unified phylogeny of all NLRs based on their shared nucleotide-binding domain (Figure 1A). This tree contained 7,818 NB-ARC sequences that had >70% coverage across the NB-ARC domain and represented 7,716 NLR genes, including 168 NB-ARC sequences of NLRs from the reference Arabidopsis Col-0 assembly. Despite the N-terminal domains not being included in the analysis, this phylogeny clearly split into clades corresponding to the three canonical architectures: RPW8, Coiled-coil, and TIR domain-containing NLRs supporting previous observations of monophyletic origin of TNLs, CNLs and RNLs (Shao et al., 2016; Tamborski and Krasileva, 2020).

**Figure 1.**
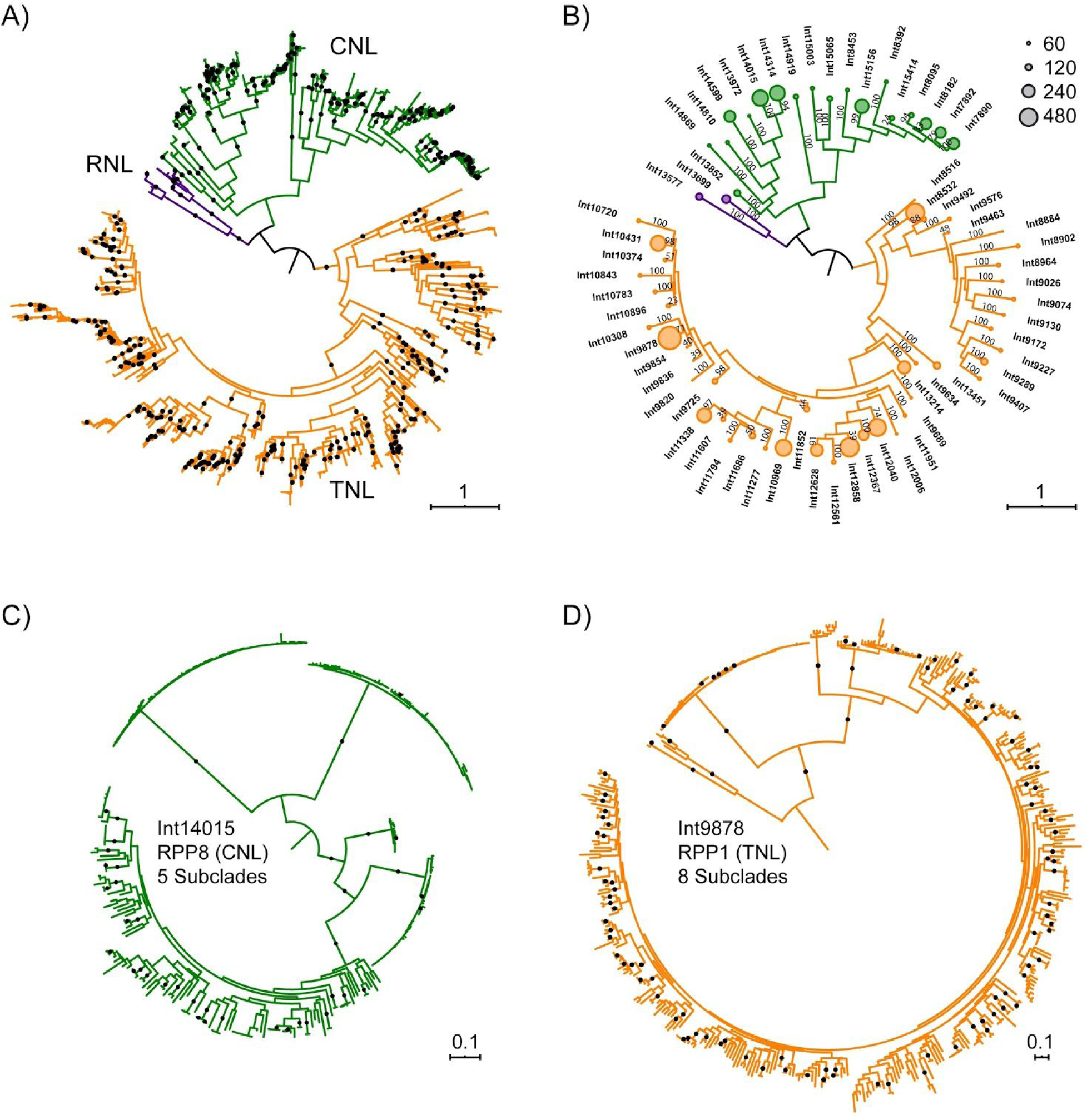
Phylogenetic analyses of Arabidopsis pan-NLRome. **A)** Maximum likelihood tree for 7,818 Arabidopsis NB-ARC sequences rooted on a branch connecting TNL and non-TNL clades. 99% or better bootstrap values shown as dots. **B)** Same tree as in A) partitioned into 65 initial clades with clade size shown as circles and indicating bootstrap support for each clade. **C)** Int14015 clade tree (rooted midpoint) based on a full-length alignment of the clade sequences. 99% or better bootstrap values shown as dots. **D)** Int9878 clade ML tree (rooted midpoint) based on a full-length alignment of the clade sequences. 99% or better bootstrap values shown as dots.

To facilitate downstream analyses, we first split the overall phylogeny into 65 cwlades, based on clade size (40-500 sequences) and bootstrap support. Of these, 43 clades had bootstrap scores of 100, 12 clades >70, and only 10 clades remained with low bootstrap values, grouping sequences that could not be confidently assigned elsewhere (Figure 1B). In the initial partition, the largest clade contained 431 sequences, allowing us to construct *de novo* full-length alignments and clade phylogenies for all clades. The tree of one of the initial clades, Int14015, containing resistance gene RPP8 is representative of observed evolutionary dynamics and is shown in Figure 1C. It reveals five well supported subclades that differ in size and internal diversity, as reflected in very short internal branch lengths in four out of five subclades. The observation that closely related sequences evolve at very different rates is true not only for RPP8, but throughout the NLR family. RPP1, a well characterized NLR, which is known to directly interact with its target, ATR1, also has closely related sequences that are largely identical in different ecotypes (Figure 1D). In fact, all clades with longer branches, i.e. higher amino acid divergence, have closely related clades with paralogous genes that show very little variation between ecotypes. These observations are consistent with NLR genes experiencing rapid shifts in the selection pressure (Ding et al., 2007).

We iteratively refined initial clades by splitting them into two or more subclades and repeating alignment and phylogeny generation steps. We prioritised cutting long, well supported internal branches and so tended to preserve both rapidly evolving and low variability subclades (see methods). After two iterations, NLRs fell into 223 non-singleton and 14 singleton clades. This NLRome partition is somewhat more conservative than the OrthoMCL-based analysis which produced 464 orthogroups and 1663 singletons (Van de Weyer et al., 2019). In our final clade assignments, 83% of all clades contained no more than one gene for all represented ecotypes, thus approximating allelic series. Over 90% of all NLRs fell into clades of 20 or more genes, allowing sampling for sequence diversity analysis. Only six large clades that ranged in size from 73 to 323 sequences contained multiple genes for ten or more ecotypes and could not be split further due to lack of long internal branches with strong support (Table S1). The large clades contained RPP1, RPP4/5, RPP39, and RPP8, suggesting that interallelic exchange complicated the phylogeny and prevented separation into allelic series. Taken together, our analyses suggest that pan-genomic NLR repertoires can be clustered into near-allelic series using phylogenetic approaches.

### Sequence analysis of the NLRome clades identifies highly variable NLRs

NLR genes code for immune receptors that provide protection during pathogen infection. Their highly variable regions are expected to contain the specificity-determining residues. We used Shannon entropy as a sensitive and robust measure of amino acid diversity. Entropy is zero at positions that are invariant and it reaches a theoretical maximum of log_2_20 or ∼4.32 when all 20 amino acids are present in equal ratios; a position with two variant amino acids present at equal ratios produces a value of 1 bit. A Shannon entropy plot thus represents a fingerprint of sequence diversity encoded in the alignment (Figure 2A).

**Figure 2.**
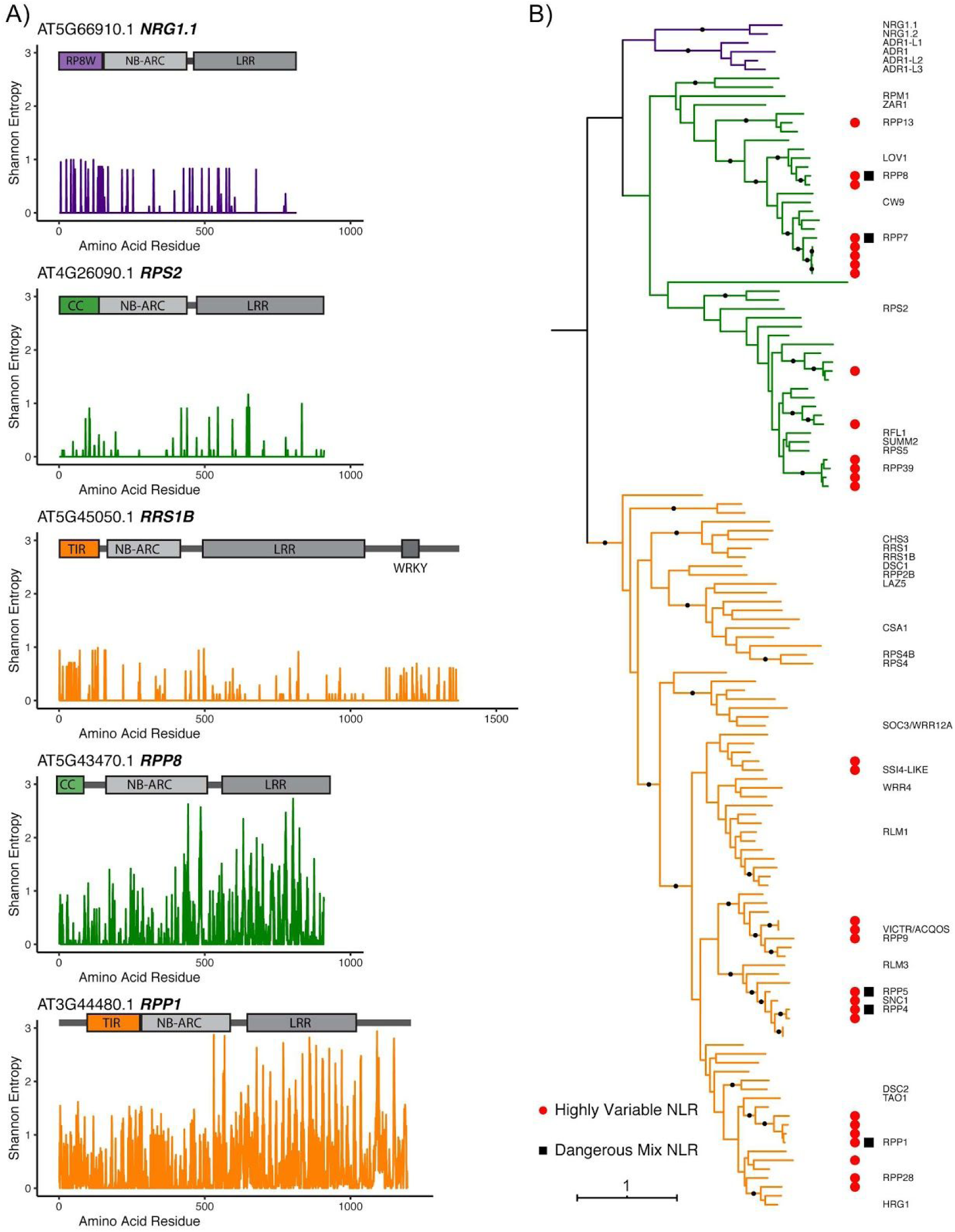
Identification and phylogenetic distribution of highly variable NLRs. **A)** Domain diagrams and Shannon entropy plots of clade alignments containing known NLRs from ancient helper (NRG1.1), guard (RPS2), integrated decoy (RRS1B), and direct recognition (RPP8 and RPP1) functional groups. **B)** Phylogenetic distributions of NLRs of the reference ecotype, Col-0, indicating positions of known genes and showing the location of hvNLRs and autoimmune Dangerous Mix (DM) NLRs. 99% or better bootstrap values shown as dots.

Several functional classes of NLRs produced entropy plots with limited diversity. An ancient helper RNL, NRG1.1, an indirect recognition CNL, RPS2, and an integrated-domain TNL, RRS1B, all produced entropy plots that showed entropy never exceeding 1 bit. Low sequence variability in these clades is consistent with their conserved functions. In contrast, a total of 30 NLR genes in the reference ecotype Col-0, including 14 CNL genes and 16 TNL genes, belonged to clades whose alignments repeatedly scored above 1.5 bits and revealed a series of periodic spikes in the LRR region. Among these genes were the known direct recognition proteins from the RPP8 and RPP1 clades. Using Shannon entropy as a metric, we defined highly variable NLRs (hvNLRs) as those with 10 or more positions exceeding 1.5 bit cutoff. No protein known to indirectly recognize pathogen effector was found among hvNLRs and all known direct binders were among hvNLRs (Figure 2B). When we ran Shannon entropy analyses on the previously identified NLR orthogroups (Van de Weyer et al., 2019), we only detected 15 hvNLRs, of which 5 did not overlap with our phylogeny-based analyses (3 slightly below 1.5 bits cutoff and 2 not supported as true orthogroups by phylogeny). This suggests that phylogeny-based orthogroup assignment specifically preserved the hvNLR clades. We predict that phylogeny-based NLR clade analysis combined with Shannon entropy can be applied to non-model plants to computationally separate candidate direct binders from other NLRs based on their sequence diversity.

### Highly variable NLRs are distributed throughout the TNL and CNL clades

We observed that hvNLRs were distributed over the NLR tree of the reference accession Col-0 with representatives in both TNL and CNL major clades. Within both major clades, there were multiple hvNLR genes right next to conserved paralogs that did not show excess diversity. This is consistent with our prior observation that NLR subclades with long branches have close paralogs with limited subclade diversity. Recent duplications of hvNLRs produce local hvNLR clusters such as those near RPP7, RPP39, RPP4/5, and RPP1. NLRs found in phylogenetic proximity often also cluster physically on the Arabidopsis chromosomes. Nonetheless, genomic clustering with close paralogs is not required for an NLR to become highly variable as shown by RPP9, RPP13, and RPP28. Also presence in a physical cluster does not force a gene to become an hvNLR as shown by RLM3 in the RPP4/5 genomic cluster and CW9 in the RPP7 genomic cluster. Thus it appears that the copy number variation that is observed in the clusters is an independent process that helps create material for NLR evolution, but generation of highly variable NLRs can proceed outside of genomic clusters.

The physical proximity and phylogenetic relationships of hvNLRs and their closely related low variability paralogs suggest rapid switches in selective pressure involved in generating the apparent diversity. Since the selection is likely to correlate with the NLR function, we located the known guardian NLRs within the phylogeny. Since these are expected to maintain binding sites for conserved plant proteins, we expected them to show low entropy scores. As we have already seen for RPS2, other known guardian NLRs including RPM1, RPS5, and ZAR1 all showed low variability. They did not however form a separate clade within the phylogeny, instead, they were interspersed by hvNLRs. This phylogenetic arrangement together with the excess of both copy number variation and of amino acid diversity in the hvNLRs argue for a mechanism where hvNLRs mostly act in direct recognition mode but are infrequently able to generate indirect recognition alleles that are preserved due to their competitive advantage.

### Highly variable NLRs contain the known NLR autoimmune loci

Generating diverse receptors in the immune system carries with it a cost of autoimmune recognition. In the known Dangerous Mix gene pairs at least one and sometimes both causative alleles are NLRs (Chae et al., 2014). If our prediction that highly variable NLRs are sources of novel direct binding is correct, we would expect to find a strong overlap between hvNLRs and Dangerous Mix NLRs. Indeed, hvNLR clades contain all the known NLR Dangerous Mix genes including RPP7, RPP8, RPP4/5 and RPP1. We suspect that in the future more DM NLRs will be found that will map to other hvNLR loci. This finding also suggests that targeted resequencing of NLRs in crop species could help identify loci responsible for hybrid necrosis phenotypes that are a frequent impediment to breeding.

### Highly variable residues cluster on the surfaces of LRR domains of hvNLRs

In plant NLRs, the LRR domains are known to encode the recognition specificities. So, at first we wanted to know whether highly variable residues occur predominantly in the LRR domain. This was indeed the case for all 30 hvNLRs considered (Table 1). We noticed however that regions in the NB-ARC domain also had high entropy scores in multiple NLRs (RPP1 and RPP8 in Figure 2A). This suggests that a limited number of residues in the NB-ARC domain could participate in target binding in these receptors. .in the LRR in absence of the ligand. Many TNLs have post-LRR domains that lack the characteristic LRR pattern of residues yet are predicted to be folded and form a contiguous structure with the preceding repeats (Van Ghelder and Esmenjaud, 2016). We observed that the post-LRR domains also often contained residues with high entropy scores (RPP1 in Figure 2A). Together these data suggest that the LRR carries the majority of binding residues, while NB-ARC and post-LRR domains can also participate in ligand binding.

**Table 1.** Number and location of highly variable residues in hvNLR receptors. Number of residues in clade alignment for each hvNLR with Sannon entropy values of at least 1.5 bits, counted by domain. Majority of highly variable residues were found in the LRR domain.

| Gene ID | Gene Name | pre NB-ARC | NB-ARC | Linker | LRR | post LRR |
| --- | --- | --- | --- | --- | --- | --- |
| AT1G31540.2 | RPP9 | 0 | 0 | 0 | 23 | 11 |
| AT1G58602.1 | RPP7 | 0 | 0 | 1 | 34 | 0 |
| AT1G58807.1 |  | 1 | 0 | 0 | 29 | 1 |
| AT1G58848.1 |  | 1 | 0 | 0 | 37 | 0 |
| AT1G59124.1 |  | 1 | 0 | 0 | 17 | 0 |
| AT1G59218.1 |  | 1 | 0 | 0 | 36 | 1 |
| AT1G61180.1 |  | 2 | 7 | 0 | 35 | 0 |
| AT1G61190.1 | RPP39 | 2 | 4 | 0 | 36 | 1 |
| AT1G61300.1 |  | 2 | 7 | 0 | 35 | 0 |
| AT1G61310.1 |  | 2 | 7 | 0 | 35 | 0 |
| AT1G62630.1 |  | 0 | 4 | 0 | 23 | 2 |
| AT1G69550.1 |  | 0 | 2 | 2 | 58 | 1 |
| AT2G14080.1 | RPP28 | 1 | 0 | 1 | 18 | 5 |
| AT3G44400.1 |  | 2 | 4 | 3 | 22 | 18 |
| AT3G44480.1 | RPP1 | 3 | 6 | 3 | 35 | 17 |
| AT3G44630.1 |  | 3 | 6 | 3 | 35 | 15 |
| AT3G44670.1 |  | 4 | 4 | 3 | 30 | 19 |
| AT3G46530.1 | RPP13 | 0 | 0 | 1 | 34 | 0 |
| AT4G16860.1 | RPP4 | 3 | 5 | 5 | 45 | 1 |
| AT4G16890.1 | SNC1 | 6 | 5 | 5 | 34 | 5 |
| AT4G16920.1 |  | 7 | 5 | 5 | 51 | 3 |
| AT4G16950.1 | RPP5 | 7 | 5 | 5 | 41 | 8 |
| AT5G38350.1 |  | 0 | 3 | 1 | 13 | 6 |
| AT5G41740.1 | SSI4-LIKE | 0 | 3 | 0 | 21 | 4 |
| AT5G41750.1 |  | 0 | 2 | 1 | 19 | 2 |
| AT5G43470.1 | RPP8 | 0 | 10 | 0 | 19 | 0 |
| AT5G43740.1 |  | 0 | 2 | 2 | 27 | 0 |
| AT5G46510.1 |  | 1 | 1 | 1 | 7 | 6 |
| AT5G46520.1 | VICTR/ACQOS | 1 | 1 | 1 | 7 | 6 |
| AT5G48620.1 |  | 0 | 10 | 0 | 19 | 0 |

If the high entropy residues do indeed make up the target binding sites, we would expect to find them in one or two clusters on the receptor surfaces and to include exposed hydrophobic residues. LRR domains fold in a predictable manner that buries the conserved leucines and exposes the variable residues on the protein surface; this allows us to skip structure prediction and to approximate LRR surfaces based on repeat annotation. The concave side of LRR domains contains a beta-sheet with a regular array of surface-exposed residues, and it can be represented as a table with one line per repeat unit and the columns corresponding to variable positions in the canonical Lx_2_x_3_Lx_5_Lx_6_x_7_ repeat. In the case of ZAR1, the first known plant NLR structure, such matrix representation based on repeat annotation perfectly matches the one that is based on the experimental structure (Figure 3A). In order to test whether entropy analysis can predict NLR binding sites, we annotated LRRs for each hvNLR gene in Col-0 and mapped entropy scores onto this representation (Figure 3B for three representative examples, Figure S1 for all Col-0 hvNLRs). This analysis revealed that in all the hvNLRs, the periodic spikes in entropy signal over the LRR likely correspond to one or two surface clusters in the NLR protein (Figure 3B, Figure S1). In the first example, AT5G43740, the strongest variability signal is found in LRRs 8 through 12 and positions 3, 5, 7, and 8 of the repeat. Additional high entropy signal comes from LRR1 through LRR5 positions 8 and 10. In RPP13, the positions C-terminal to the predicted beta sheet appear to play an important role in determining binding specificity. Unlike AT5G43740, highly variable residues in positions 8, 9, and 10 of the repeats appear throughout the annotated LRR region, while all residues in position 2 and 3 are conserved. We therefore predict that in RPP13 loops that follow the beta strands play a key role in determining substrate specificity. Our prediction that specificity determinants of RPP13 stretch between LRR1 and LRR12 are in agreement with the large experimentally identified RPP13 specificity determining region (Rentel et al., 2008).

**Figure 3.**
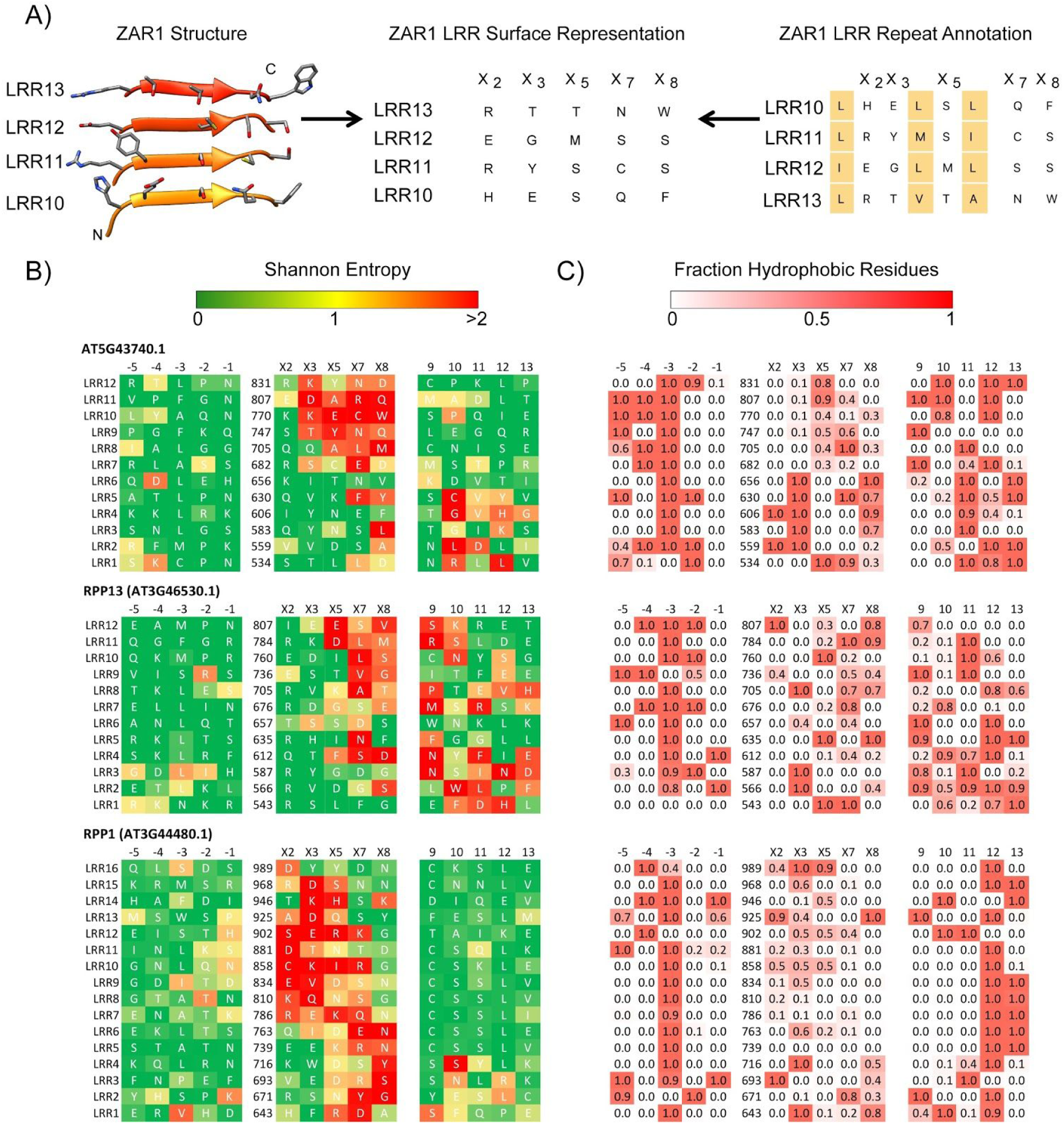
2D representations of LRR surfaces allow comparisons of predicted NLR binding sites in absence of experimental structures. **A)** Beta-sheet on the concave side of ZAR1 LRR domain shows a regular array of surface-exposed residues that correspond to the variable positions in the LxxLxLxx LRR motif (left). Single-letter amino acid representation of the observed array (center). Identical representation is obtained from LRR repeat annotation by arranging the rows from bottom to top and hiding the columns containing conserved leucines (right). **B)** Shannon entropy scores and amino acids of three representative Col-0 hvNLRs mapped onto the 2D surface representation including additional 5 amino acids on either side of the core repeat unit. **C)** Percent hydrophobic residues in the alignments of the same three proteins.

RPP1 is a well studied example of direct recognition NLR where multiple alleles have different recognition profiles with regard to the alleles of the *Hyaloperonospora arabidopsis* effector ATR1 (Rehmany et al., 2005). In RPP1, we observed a large number of contiguous residues that likely contribute to binding specificity stretching from LRR1 to LRR15. Highly variable residues are concentrated in positions 5, 7 and 8 at the beginning of the domain, but shift towards the start of the beta strands in the later repeat units with residues 2, 3 and 5 lighting up uniformly in LRR7 - LRR15. Rather unusually, we also observe some variable residues in −1 and −2 positions. We conclude that in RPP1 (and in AT5G43740), the targets likely bind through the middle of the horseshoe LRR shape, rather than on one side of it as in the case of RPP13. The predicted RPP1 binding site contains the residues previously shown to extend recognition specificity of RPP1 allele NdA towards ATR1-Maks9 (Krasileva, 2011).

To test the prediction that the identified highly variable surfaces represent target-binding sites, we surveyed these regions of high diversity for presence of exposed hydrophobic residues, which are commonly found at the centers of protein-protein binding sites (Figure 3C). Indeed, in every case the highly variable residues included exposed hydrophobic amino acids often including bulky aromatics such as tryptophan and phenylalanine. We also tested whether the entropy based predictions agree with positive selection analyses that have been used in the past to identify functionally important residues in NLRs (Kuang et al., 2004). In RPP13 66% of all high-entropy residues (>1.5 bits) were under positive selection according to Phylogenetic Analysis by Maximum Likelihood (PAML) Model 8 (Figure S2). All of the remaining high-entropy residues fell into regions that contained gaps in the alignment and could not be analysed by PAML. Thus, entropy analyses of hvNLR surfaces can help identify functionally important residues that likely determine receptor binding specificity.

### ZAR1-RKS1 binding site overlaps RPP13 entropy-based prediction

Arabidopsis ZAR1 is an indirect-recognition NLR and the only one to date with a known structure. In our phylogeny, it is closely related to the hvNLR RPP13 (Figure 2B). While the ZAR1 entropy plot lacked high entropy residues, we wanted to compare the known footprint of RKS1, the ZAR1 binding partner, with the positions of highly variable residues in RPP13. Unusually for hvNLRs, RPP13 highly variable residues cluster on the C-terminal side of the repeats, with positions 7-10 of the repeat units showing the highest diversity (Figure 3B). Surprisingly, the similarly positioned residues in ZAR1 are used to bind its stable complex partner, RKS1 (Figure 4). This finding is consistent with ZAR1 and RPP13 emerging from an hvNLR common ancestor that had a binding site similar to that observed in ZAR1 and predicted in RPP13.

**Figure 4.**
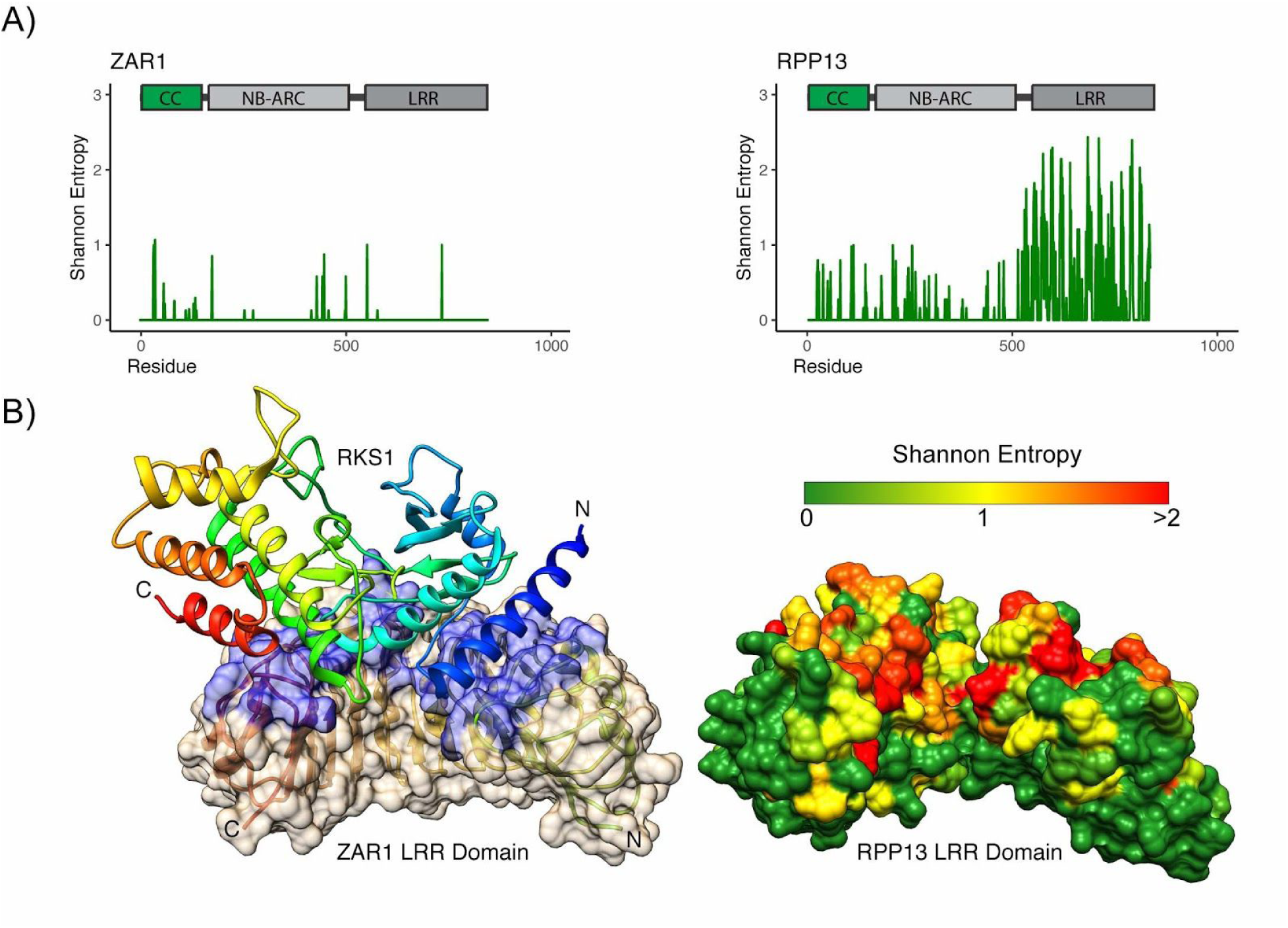
RPP13 predicted ligand binding site overlaps observed ZAR1-RKS1 binding site. **A)** Shannon entropy plots and domain diagrams for ZAR1, an indirect recognition CNL, and RPP13, a related hvNLR. **B)** Cryo-EM structure of RKS1 bound to ZAR1 (CC and NB-ARC domains omitted for clarity) (PDB ID: 6J5W). RKS1 shown as a secondary structure diagram with rainbow coloring from blue (N-terminus) to red (C-terminus), ZAR1 LRR as secondary structure diagram and transparent surface with RKS1 contact residues colored blue. RPP13 LRR domain homology model as surface oriented as ZAR1 and colored by Shannon Entropy of the RPP13 clade alignment from low (green) to high entropy (red).

**Figure 5.**
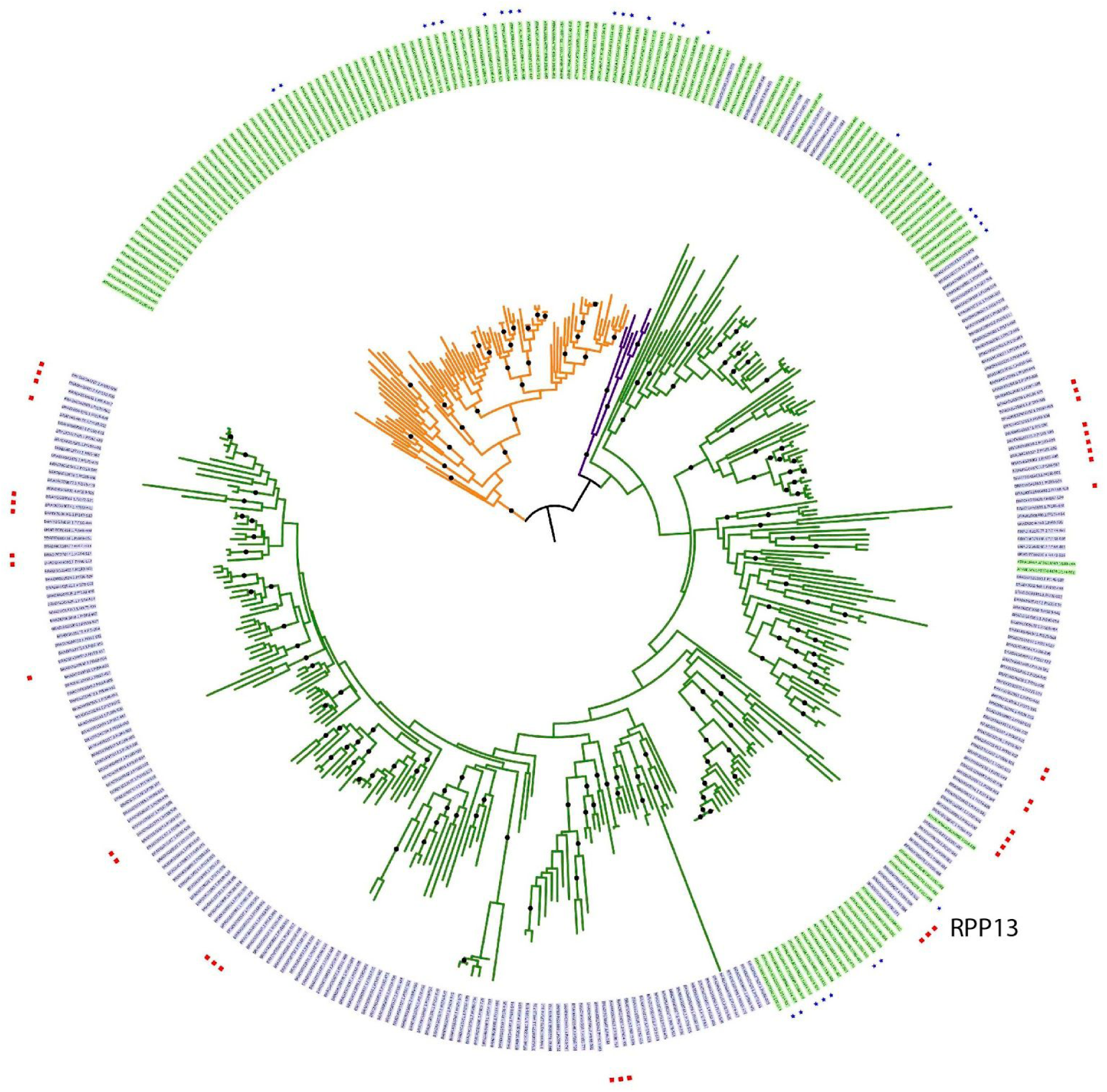
Dispersed distribution of hvNLRs in a joint phylogeny of Brachypodium Bd21 (blue labels) and Arabidopsis Clo-0 (green labels). The Arabidosis hvNLR clades (red dots) and Brachypodium hvNLRs (blue dots) do not cluster except for the RPP13 CNL clades. The tree is rooted arbitrarily on a branch connecting TNL clade (orange branches) and non-TNL clades (RNL branches are in purple and CNL in green). 99% or better bootstrap values shown as dots.

### Similar phylogenetic distribution of hvNLRs in Brachypodium distachyon

In order to test whether our methods and findings were applicable beyond A. thaliana, we performed a similar analysis on 54 lines of Brachypodium distachyon, a model grass species. The automatic short-read assembly and annotation pipeline used in generating the Brachypodium data is less reliable than the targeted resequencing approach used to generate Arabidopsis pan-NLRome. Specifically, only 45% of hvNLRs present in the reference strain, Bd-21, were recovered in the assembly control. Nonetheless, the overall picture that emerges from the analysis of Brachypodium NLR clades is similar to that of Arabidopsis. After splitting the overall Brachypodium NLR tree into 91 initial clades, we performed 4 rounds of clade refinement to arrive at a final clade partition with 433 subclades. Of these, 28 produced alignments that fulfilled the hvNLR criteria. Altogether, 40 hvNLRs in the reference accession, Bd21, were identified as hvNLRs.

Similar to A. thaliana, Brachypodium hvNLRs were distributed throughout the phylogeny including in the highly expanded monocot-specific CNL clade. Here too, hvNLRs had sister clades that showed little amino-acid diversity. Importantly, when we constructed the joint tree for Col-0 and Bd21 reference NLRomes, the only hvNLRs from the two species that lay close together belonged to the RPP13-like clades (Figure 4). This highlights the importance of sequencing of pan-NLRomes of plants of interest as identification of hvNLRs is unlikely to be transferable except for closely related species.

## Discussion

Even before the first NLR structure or the extensive sequence datasets were available Mitchemore and Meyers predicted that hypervariable amino acid positions in the NLRs would map to the concave surface of the LRR domain based on the signatures of positive selection in a number of selected examples (Michelmore and Meyers, 1998). They generalized that this might be true for all NLRs. This model was challenged by the discovery of indirect recognition and strongly conserved NLRs. Our analysis proposes a new methodology to study NLR-omes, predicts NLR mode of action through sequence analysis, and reconciles evolution of direct recognition NLRs (under diversifying selection) and indirect recognition NLRs (under purifying or balancing selection).

In this paper, we observed that hvNLRs account for the known direct recognition NLRs and for autoimmune NLRs. We also observed that the hvNLRs have close paralogs with little allelic diversity that include the known indirect recognition NLRs. Based on this observation, we propose that indirect recognition NLRs are a functional byproduct of hvNLR evolution providing an important update of the birth-and-death model (Michelmore and Meyers, 1998). We conclude that continuous diversity generation acts on a limited subset of NLR genes creating a wide recognition potential, including binding to endogenous plant proteins. When recognition of endogenous proteins is beneficial, such as under perturbations by the pathogen, it evolves into indirect recognition and starts to experience different selective forces.

The resolution and sensitivity of our analyses became possible by adopting two key approaches: identification of orthologous groups of NLR receptors by phylogeny in place of commonly used distance metrics and using simpler Shannon entropy measure of diversity in place of more complex evolutionary models. Separating rapidly evolving protein families into meaningful clades or groups for downstream analyses is a common challenge. In the NLR family of plant immune receptors, it is further complicated by ongoing information flow between close paralogs through recombination and gene conversion (Kuang et al., 2004). Phylogeny-based analyses are considered to be more accurate than distance based methods in similar problems such as classification of Human Immunodeficiency Virus isolates (Pineda-Peña et al., 2013). Our phylogeny-based partition of NLR immune receptors into clades improved on the published OrthoMCL-based partition by producing more encompassing clades and in particular fewer singletons. OrthoMCL is a distance-based algorithm that was originally developed to separate members of different protein families rapidly; it uses a single parameter to determine the rate of convergence (Li et al., 2003). This makes its use to partition the pan-NLRome problematic, because closely related NLRs are known to experience vastly different selection pressures and thus are expected to contain very different amounts of allelic diversity (Bakker et al., 2006; Kuang et al., 2004). The specific danger for hvNLR identification is that highly variable clades will be split, losing the relevant signal. This is indeed what we observed, as the OrthoMCL-based analysis identified only one out of three hvNLRs and missed key sources of new NLR specificity such as the RPP1 cluster, which got split into small orthogroups. The drawback of the phylogeny-based approach is that it is not yet fully automated, however we are hopeful that phylogeny-aware algorithms will emerge that will fill this gap. One alternate approach that would simplify the analysis would be to replace the initial clade assignment with iterative matching of NLR sequences against a set of inferred ancestral NLR models (Shao et al., 2016).

It has been well established that closely related NLRs experience different modes of selection (Ding et al., 2007; Wang et al., 2011; Kuang et al., 2004). By expanding this observation to the pan-NLRome and combining it with the wealth of characterized NLRs in Arabidopsis, we were able to decipher a larger evolutionary pattern where hvNLRs act as sources of new specificities and encompass the known direct-recognition NLRs. Their diversification, while advantageous to the plant, comes at a cost. All known Dangerous Mix NLR genes that can trigger autoimmune recognition belong to hvNLR clades. Thus, generation of novel specificities goes hand in hand with potential for self-recognition and auto-immunity. We also propose that during their continuous evolution, hvNLRs can generate indirect-recognition NLRs at a low frequency. Because indirect recognition usually tracks a conserved effector activity, it is more robust than direct recognition of the effector surface. Duplication of such successful variants might then be favored due to the increased fitness of the progeny where one copy could eventually be preserved while the other could continue to generate novel specificities (Kondrashov et al., 2002). The latter inference is consistent with our observation that ZAR1, an indirect-recognition NLR, binds to its stable complex partner, RKS1, through the same surface on the LRR that contains highly variable residues in RPP13, its closest hvNLR.

When we applied Shannon entropy analysis to NLR clades, we saw that only a subset of clades gave strong signals and that these clades included known direct recognition NLRs and autoimmune NLRs. When we looked at the distribution of high-entropy amino acids in the 30 hvNLRs of Arabidopsis reference strain, Col-0, we found that these residues commonly clustered on the predicted surfaces of LRR domains. This observation is consistent with multiple genetic and biochemical studies that support that binding specificities are largely encoded in the LRR domains (Ellis et al., 2007; Krasileva et al., 2010) as well as evolutionary studies that predict amino acid residues under positive selection to be located within LRRs (Kuang et al., 2004; Rose et al., 2004; Wang et al., 2011). When we carried out a positive selection analysis on the RPP13 clade, we found that the majority of residues with entropy >1.5 bits were under positive selection. The only exceptions were residues that could not be analysed for positive selection due to presence of gaps in the relevant alignment columns. Shannon entropy calculation does not count gap characters. It works without making complex assumptions about the data and is therefore much faster computationally.

In our analysis, we went a step further to predict binding sites in hvNLRs directly from pan-NLRome sequence data. As identified, the binding sites are large, this is likely in part due to the concave shape of the LRR scaffold that can put many of the beta strands in contact with a relatively small target. Comparisons of antibody sequence-based predictions with experimental structures shows that predictions correctly recover ∼80% of residues that do contact the antigen, while also producing many false-positives (<50% precision) (Kunik et al., 2012). Unlike the antibody, where the binding determinants are present on loops, away from the core of the structure, in the LRR a lot of predicted binding residues fall within the beta sheet located on the concave side of the domain. This suggests that the accuracy of the prediction might be higher in this system because of stronger structural constraints. Additional highly variable residues were located in post-LRR domain and in specific sites within NB-ARC, suggesting their involvement in substrate binding, or in case of NB-ARC of a compensatory mechanism to maintain self inhibition in absence of the ligand. Further mutational and structural experiments in well established NLR-effector systems would be needed to test the accuracy of these predictions and to help refine them.

Identification of the immense allelic diversity across hvNLRs argues that plant immunity is not far in its allele generation potential from the most well known adaptive immune systems. Indeed, LRRs are deployed in the adaptive immune system of lower vertebrates demonstrating that their modularity is sufficient for generation of binding to any foreign molecule (Das et al., 2013; Han et al., 2008). In the case of plants, enormous diversity is generated on population level rather than within a single organism, and therefore defending against new pathogens is a community effort. Identification of specific genes within crop species capable of such diversity generation, and their deployment in protein engineering efforts can provide valuable material for plant health.

We conclude that phylogenetic analysis of pan-NLRomes combined with Shannon entropy can rapidly classify NLRs into functional groups given sequencing information for at least 40-60 diverse samples. We also believe that our analyses would be generally applicable to identification of highly variable receptor-like proteins, such as Cf-9 in tomato (Wulff et al. 2009), and prediction of binding sites of highly variable extracellular immune receptors. Our analyses can also predict incompatibility loci which can be taken into account in breeding new crop varieties. Similar allelic diversity analyses in other non-vertebrate eukaryotes with expanded immune receptor families are needed to test whether the patterns of innate immune receptor evolution we observed are shared across the eukaryotic kingdoms of life.

## Acknowledgements

We thank members of the Krasileva lab and of the Berkeley Lab Advanced Light Source structural biology community for helpful discussions. We thank Brian Staskawicz, Raoul Martin, and Kyungyong Seong for their advice and for critical reading of the manuscript. We thank Marc Allaire and members of the Berkeley Center for Structural Biology for support, encouragement, and the use of computational resources.

---

**Supplementary Table 1.**
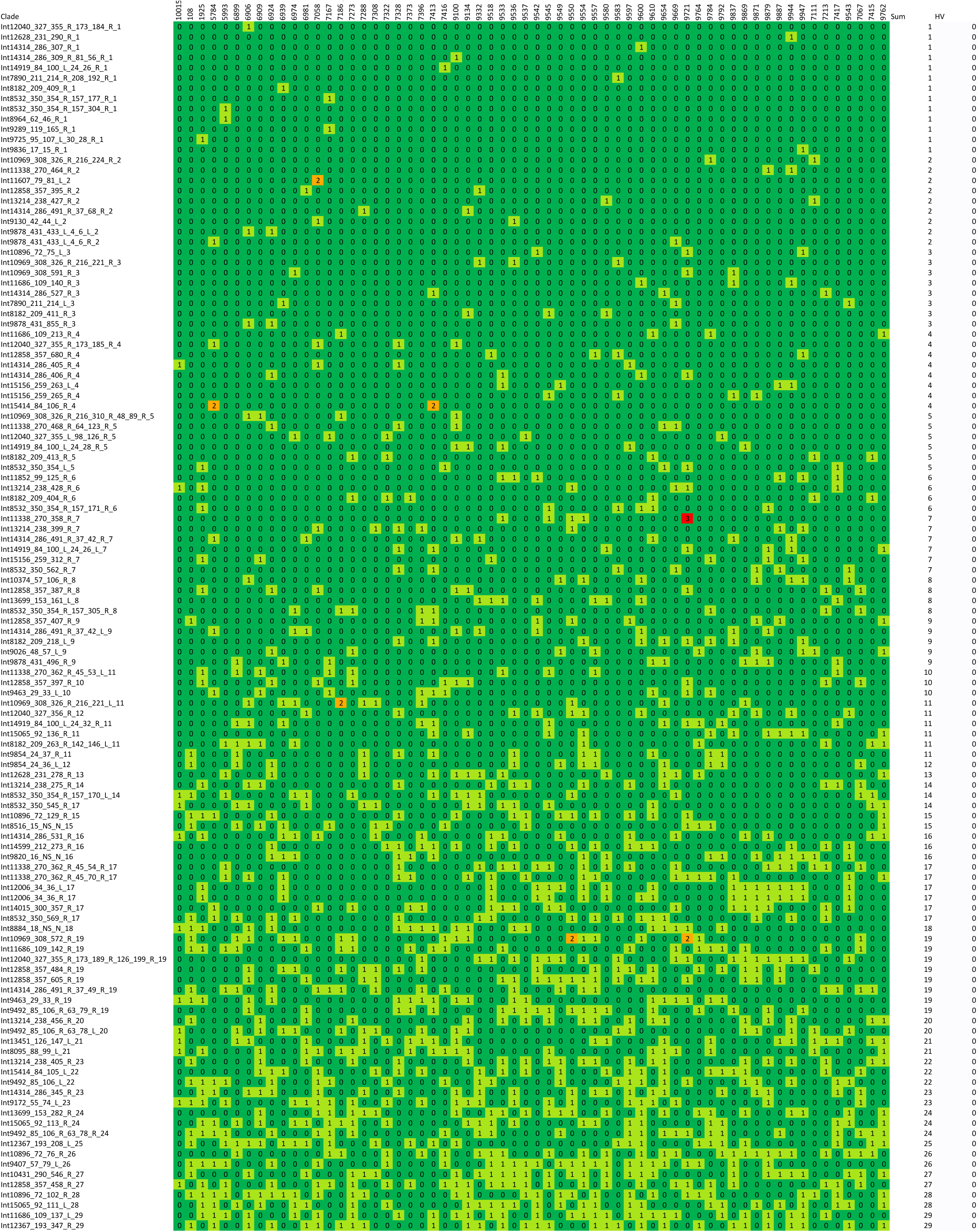
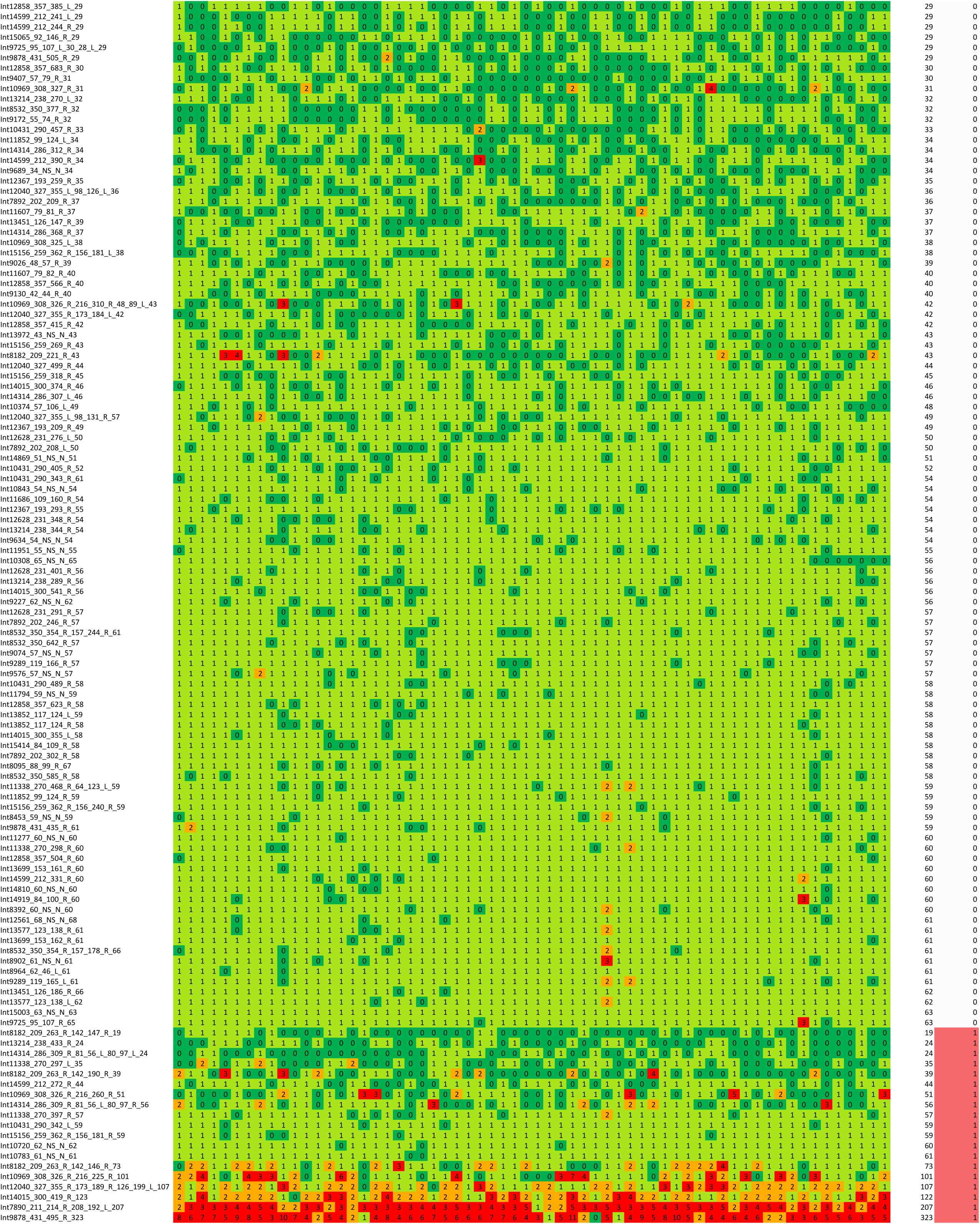
Clade representation in the 62 Arabidopsis thaliana ecotypes. The HV column value of 1 indicates that the clade alignment satisfies the hvNLR requirement.

**Supplementary Figure 1.**
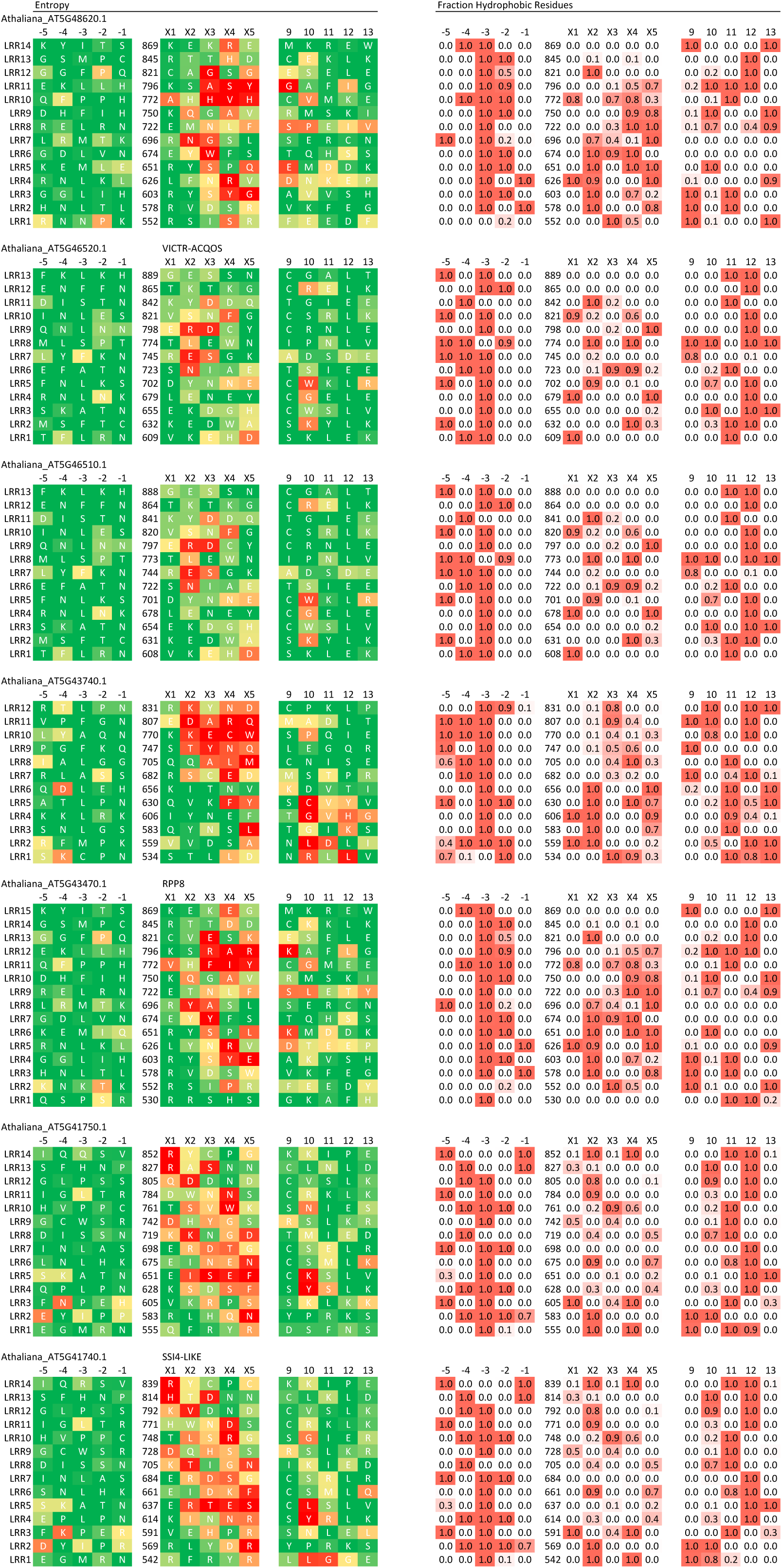
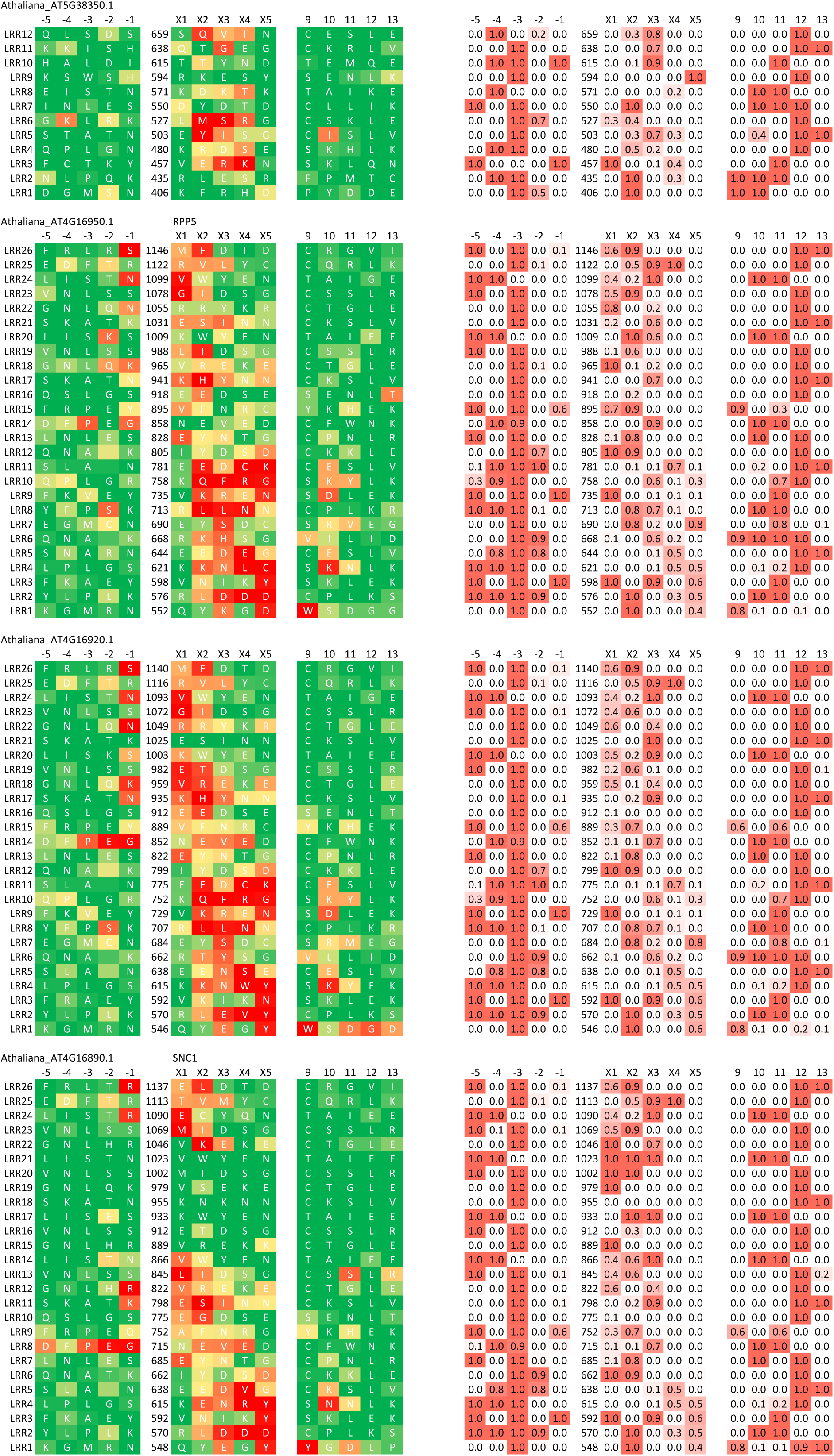
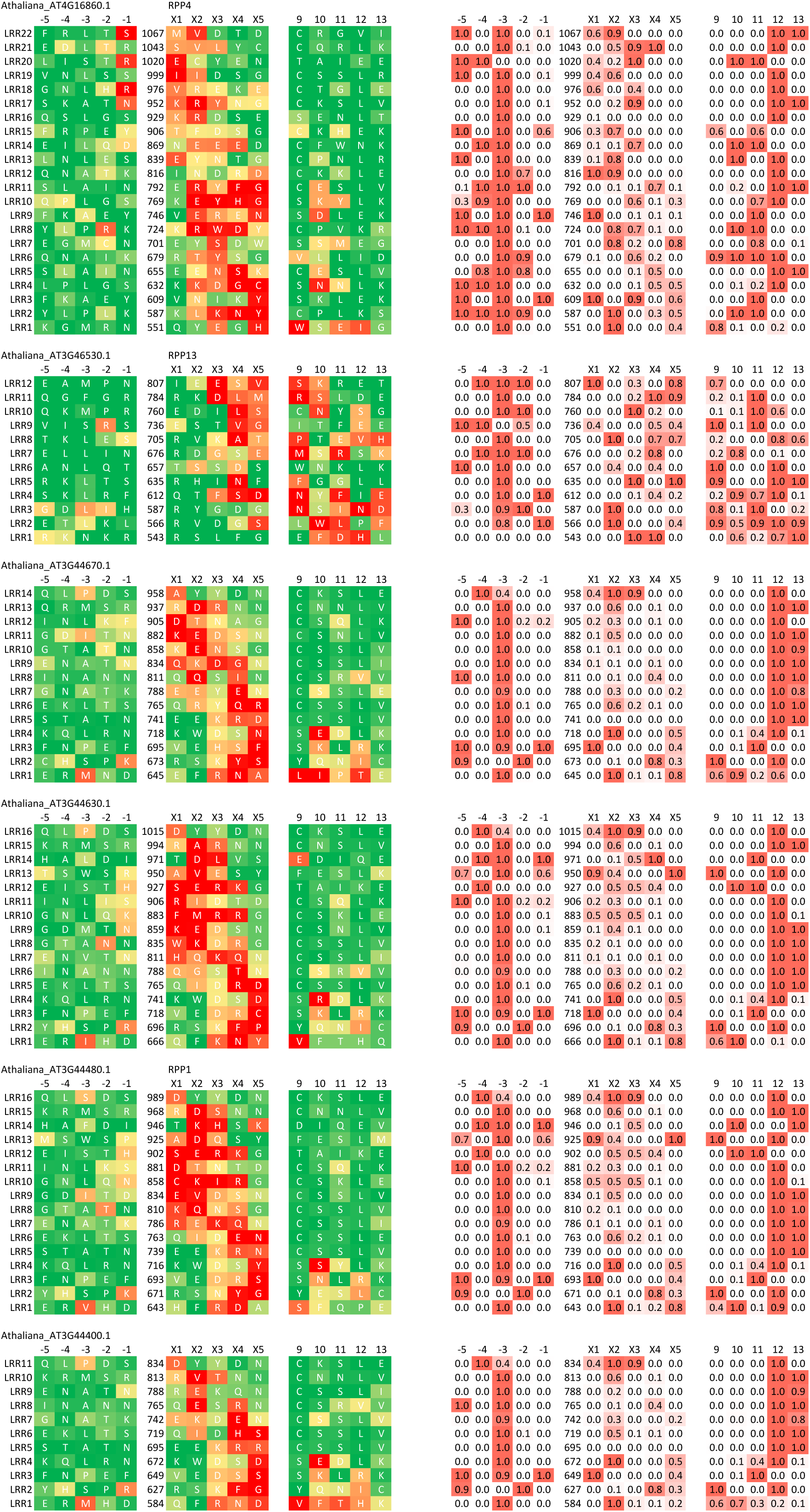
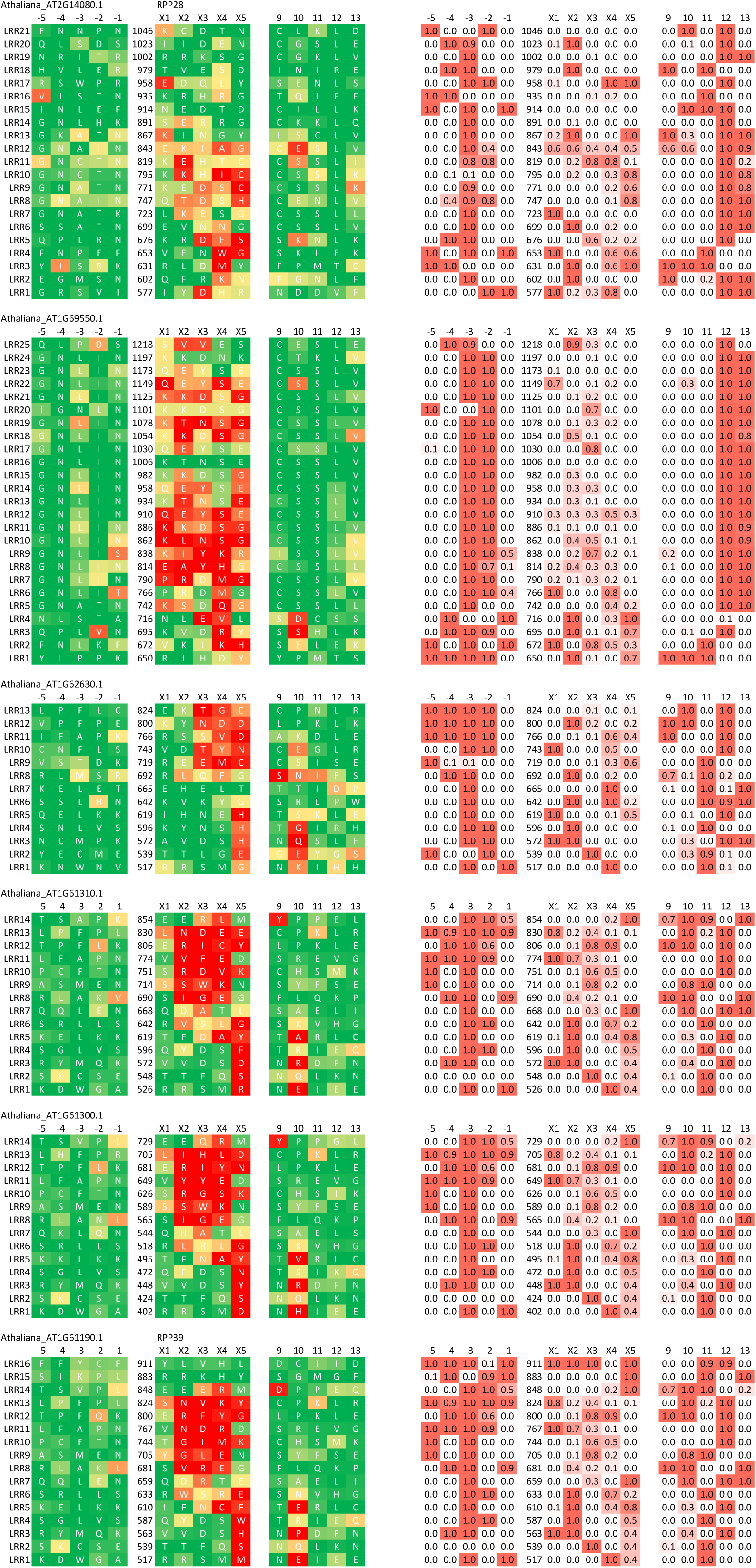
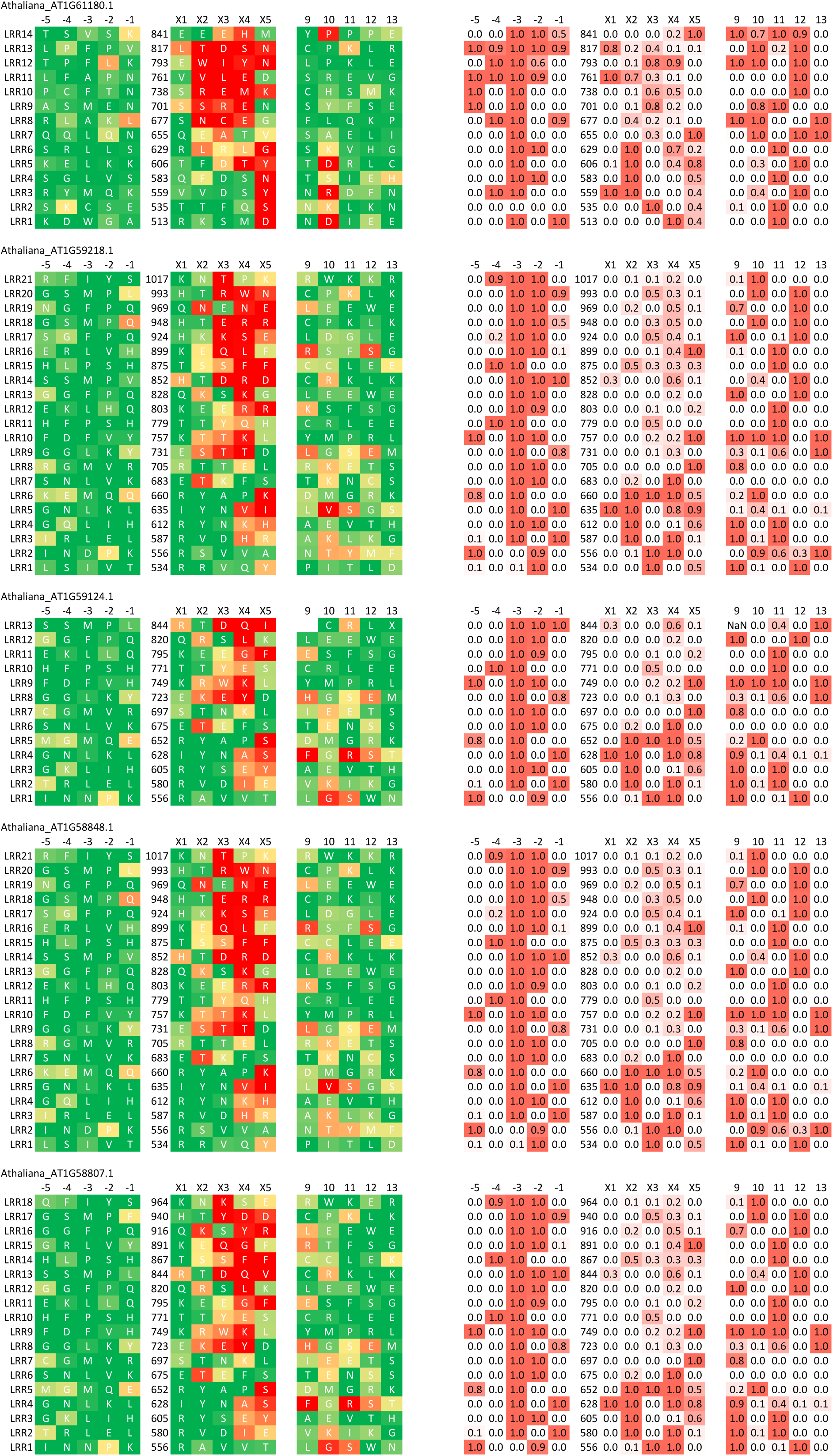
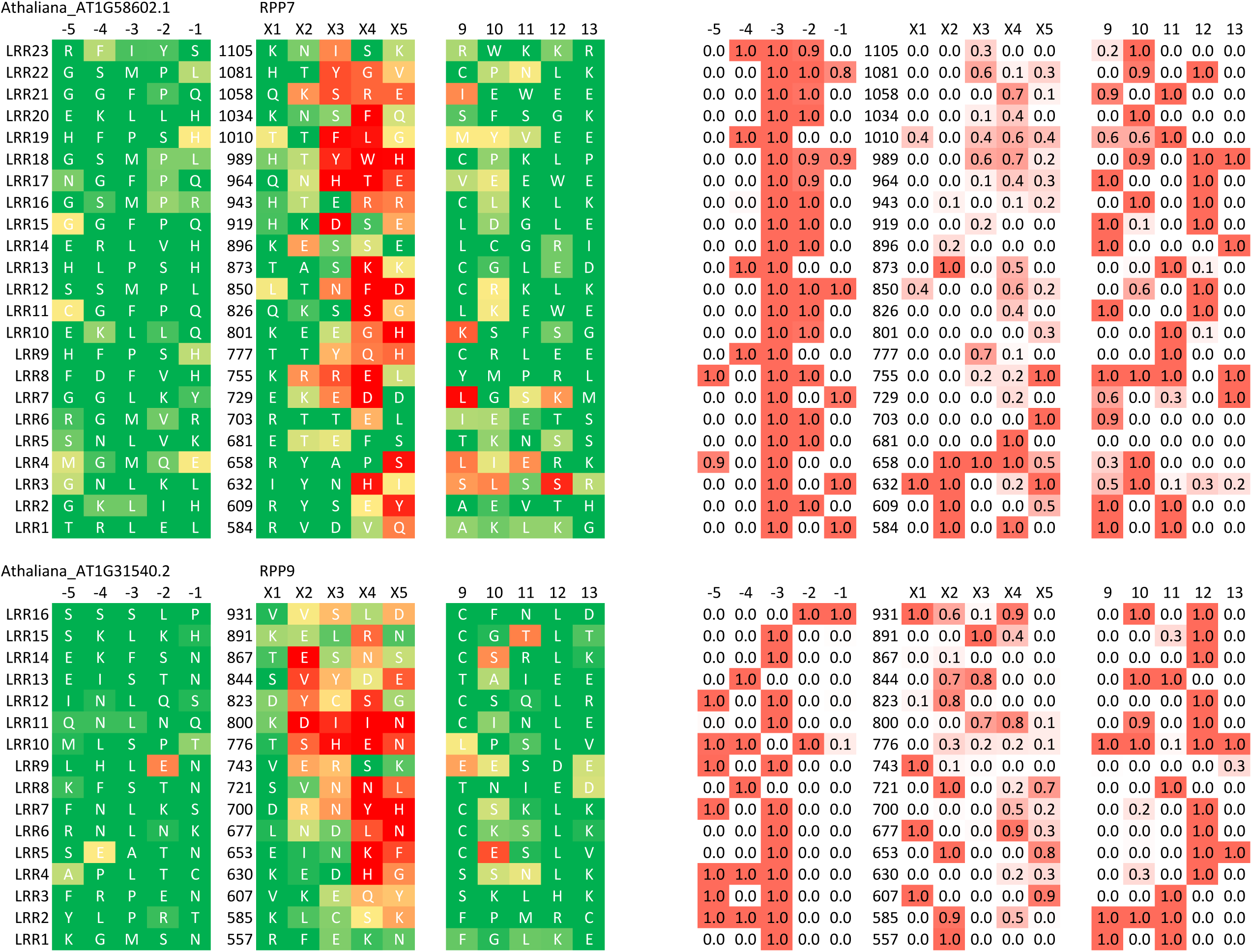
2D representations of LRR surfaces of 30 hvNLRs from ecotype Col-0.

**Supplemental Figure 2.**
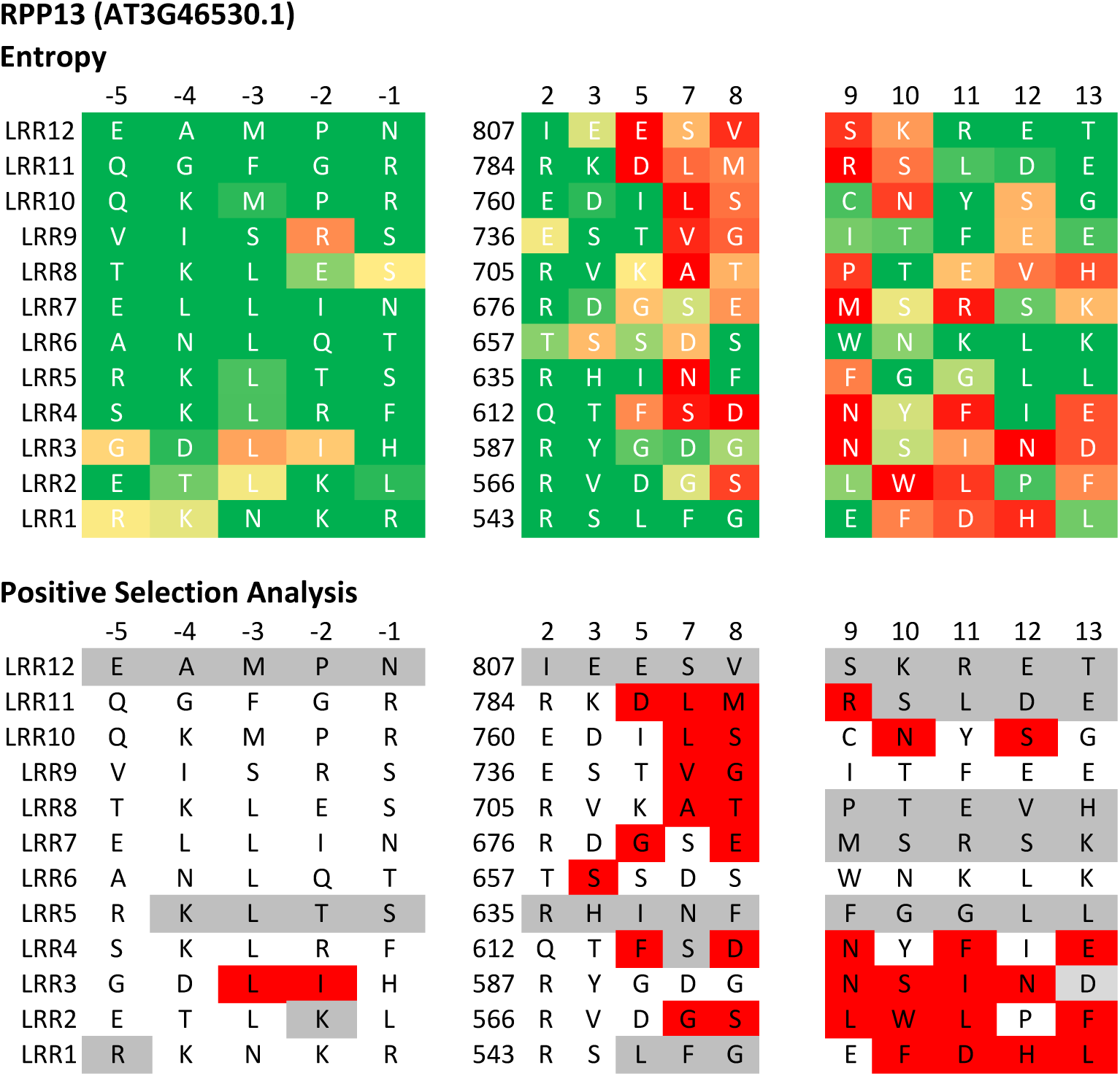
Comparison of entropy-based and positive selection-based binding site predictions. 2D surface representation of hvNLR RPP13 showing that high entropy residues are likely under positive selection. Top panel colored as in Figure 3A from low entropy (green) to high entropy (red). Bottom panel highlights in red the residues under positive selection identified with paml (>95% posterior probability). Grey background residues were not analyzed due to presence of gaps in the alignment.

## Notes

### Competing Interest Statement

The authors have declared no competing interest.

## References

Atanasov, K.E., Liu, C., Erban, A., Kopka, J., Parker, J.E., and Alcázar, R. (2018). Mutations Suppressing Immune Hybrid Incompatibility and Their Effects on Disease Resistance. Plant Physiol. 177 : 1152–1169.

Baggs, E., Dagdas, G., and Krasileva, K.V. (2017). NLR diversity, helpers and integrated domains: making sense of the NLR IDentity. Curr. Opin. Plant Biol. 38 : 59–67.

Bailey, P.C., Schudoma, C., Jackson, W., Baggs, E., Dagdas, G., Haerty, W., Moscou, M., and Krasileva, K.V. (2018). Dominant integration locus drives continuous diversification of plant immune receptors with exogenous domain fusions. Genome Biol. 19 : 23.

Bakker, E.G., Toomajian, C., Kreitman, M., and Bergelson, J. (2006). A genome-wide survey of R gene polymorphisms in Arabidopsis. Plant Cell 18 : 1803–1818.

Bomblies, K. (2009). Too much of a good thing? Hybrid necrosis as a by-product of plant immune system diversification. Botany 87 : 1013–1022.

Bomblies, K., Lempe, J., Epple, P., Warthmann, N., Lanz, C., Dangl, J.L., and Weigel, D. (2007). Autoimmune response as a mechanism for a Dobzhansky-Muller-type incompatibility syndrome in plants. PLoS Biol. 5 : e236.

Catanzariti, A.-M., Dodds, P.N., Ve, T., Kobe, B., Ellis, J.G., and Staskawicz, B.J. (2010). The AvrM Effector from Flax Rust Has a Structured C-Terminal Domain and Interacts Directly with the M Resistance Protein. Molecular Plant-Microbe Interactions 23 : 49–57.

Cesari, S. (2018). Multiple strategies for pathogen perception by plant immune receptors. New Phytol. 219 : 17–24.

Chae, E. et al. (2014). Species-wide genetic incompatibility analysis identifies immune genes as hot spots of deleterious epistasis. Cell 159 : 1341–1351.

Chen, J. et al. (2017). Loss of by somatic exchange in stem rust leads to virulence for resistance in wheat. Science 358 : 1607–1610.

Dangl, J.L., Horvath, D.M., and Staskawicz, B.J. (2013). Pivoting the plant immune system from dissection to deployment. Science 341 : 746–751.

Das, S., Hirano, M., Aghaallaei, N., Bajoghli, B., Boehm, T., and Cooper, M.D. (2013). Organization of lamprey variable lymphocyte receptor C locus and repertoire development. Proc. Natl. Acad. Sci. U. S. A. 110 : 6043–6048.

Ding, J., Cheng, H., Jin, X., Araki, H., Yang, Y., and Tian, D. (2007). Contrasting patterns of evolution between allelic groups at a single locus in Arabidopsis. Genetica 129 : 235–242.

Ellis, J.G., Dodds, P.N., and Lawrence, G.J. (2007). Flax Rust Resistance Gene Specificity is Based on Direct Resistance-Avirulence Protein Interactions. Annual Review of Phytopathology 45 : 289–306.

Gordon, S.P. et al. (2017). Extensive gene content variation in the Brachypodium distachyon pan-genome correlates with population structure. Nat. Commun. 8 : 2184.

Goritschnig, S., Steinbrenner, A.D., Grunwald, D.J., and Staskawicz, B.J. (2016). Structurally distinct Arabidopsis thaliana NLR immune receptors recognize tandem WY domains of an oomycete effector. New Phytol. 210 : 984–996.

Han, B.W., Herrin, B.R., Cooper, M.D., and Wilson, I.A. (2008). Antigen recognition by variable lymphocyte receptors. Science 321 : 1834–1837.

Jones, J.D.G., Vance, R.E., and Dangl, J.L. (2016). Intracellular innate immune surveillance devices in plants and animals. Science 354.

Jubic, L.M., Saile, S., Furzer, O.J., El Kasmi, F., and Dangl, J.L. (2019). Help wanted: helper NLRs and plant immune responses. Curr. Opin. Plant Biol. 50 : 82–94.

Kabat, E.A. (1970). Heterogeneity and structure of antibody-combining sites. Ann. N. Y. Acad. Sci. 169 : 43–54.

Kelley, L.A., Mezulis, S., Yates, C.M., Wass, M.N., and Sternberg, M.J.E. (2015). The Phyre2 web portal for protein modeling, prediction and analysis. Nat. Protoc. 10 : 845–858.

Kondrashov, F.A., Rogozin, I.B., Wolf, Y.I., and Koonin, E.V. (2002). Selection in the evolution of gene duplications. Genome Biol. 3 : RESEARCH0008.

Kozlov, A.M., Darriba, D., Flouri, T., Morel, B., and Stamatakis, A. (2019). RAxML-NG: a fast, scalable and user-friendly tool for maximum likelihood phylogenetic inference. Bioinformatics 35 : 4453–4455.

Krasileva, K.V. (2011). The Molecular Basis for Recognition of Oomycete Effectors in Arabidopsis. Doctoral Dissertation. UC Berkeley. https://escholarship.org/uc/item/57x979rf

Krasileva, K.V., Dahlbeck, D., and Staskawicz, B.J. (2010). Activation of an Arabidopsis resistance protein is specified by the in planta association of its leucine-rich repeat domain with the cognate oomycete effector. Plant Cell 22 : 2444–2458.

Kruger, J. (2002). A Tomato Cysteine Protease Required for Cf-2-Dependent Disease Resistance and Suppression of Autonecrosis. Science 296 : 744–747.

Kuang, H., Woo, S.-S., Meyers, B.C., Nevo, E., and Michelmore, R.W. (2004). Multiple genetic processes result in heterogeneous rates of evolution within the major cluster disease resistance genes in lettuce. Plant Cell 16 : 2870–2894.

Kunik, V., Peters, B., and Ofran, Y. (2012). Structural consensus among antibodies defines the antigen binding site. PLoS Comput. Biol. 8 : e1002388.

Letunic, I. and Bork, P. (2019). Interactive Tree Of Life (iTOL) v4: recent updates and new developments. Nucleic Acids Res. 47 : W256–W259.

Liao, H., Yeh, W., Chiang, D., Jernigan, R.L., and Lustig, B. (2005). Protein sequence entropy is closely related to packing density and hydrophobicity. Protein Eng. Des. Sel. 18 : 59–64.

Li, L., Stoeckert, C.J., Jr, and Roos, D.S. (2003). OrthoMCL: identification of ortholog groups for eukaryotic genomes. Genome Res. 13 : 2178–2189.

Löytynoja, A. (2014). Phylogeny-aware alignment with PRANK. Methods Mol. Biol. 1079 : 155–170.

Magliery, T.J. and Regan, L. (2005). 10.1186/1471-2105-6-240. BMC Bioinformatics 6 : 240.

Martin, E.C., Sukarta, O.C.A., Spiridon, L., Grigore, L.G., Constantinescu, V., Tacutu, R., Goverse, A., and Petrescu, A.-J. (2020). LRRpredictor-A New LRR Motif Detection Method for Irregular Motifs of Plant NLR Proteins Using an Ensemble of Classifiers. Genes 11.

Michelmore, R.W. and Meyers, B.C. (1998). Clusters of resistance genes in plants evolve by divergent selection and a birth-and-death process. Genome Res. 8 : 1113–1130.

Mistry, J., Finn, R.D., Eddy, S.R., Bateman, A., and Punta, M. (2013). Challenges in homology search: HMMER3 and convergent evolution of coiled-coil regions. Nucleic Acids Res. 41 : e121.

Pettersen, E.F., Goddard, T.D., Huang, C.C., Couch, G.S., Greenblatt, D.M., Meng, E.C., and Ferrin, T.E. (2004). UCSF Chimera--a visualization system for exploratory research and analysis. J. Comput. Chem. 25 : 1605–1612.

Pineda-Peña, A.-C., Faria, N.R., Imbrechts, S., Libin, P., Abecasis, A.B., Deforche, K., Gómez-López, A., Camacho, R.J., de Oliveira, T., and Vandamme, A.-M. (2013). Automated subtyping of HIV-1 genetic sequences for clinical and surveillance purposes: performance evaluation of the new REGA version 3 and seven other tools. Infect. Genet. Evol. 19 : 337–348.

Quinlan, A.R. (2014). BEDTools: The Swiss-Army Tool for Genome Feature Analysis. Curr. Protoc. Bioinformatics 47 : 11.12.1–34.

Rehmany, A.P., Gordon, A., Rose, L.E., Allen, R.L., Armstrong, M.R., Whisson, S.C., Kamoun, S., Tyler, B.M., Birch, P.R.J., and Beynon, J.L. (2005). Differential recognition of highly divergent downy mildew avirulence gene alleles by RPP1 resistance genes from two Arabidopsis lines. Plant Cell 17 : 1839–1850.

Rentel, M.C., Leonelli, L., Dahlbeck, D., Zhao, B., and Staskawicz, B.J. (2008). Recognition of the Hyaloperonospora parasitica effector ATR13 triggers resistance against oomycete, bacterial, and viral pathogens. Proc. Natl. Acad. Sci. U. S. A. 105 : 1091–1096.

Rose, L.E., Bittner-Eddy, P.D., Langley, C.H., Holub, E.B., Michelmore, R.W., and Beynon, J.L. (2004). The maintenance of extreme amino acid diversity at the disease resistance gene, RPP13, in Arabidopsis thaliana. Genetics 166 : 1517–1527.

Sanders, M.P.A., Fleuren, W.W.M., Verhoeven, S., van den Beld, S., Alkema, W., de Vlieg, J., and Klomp, J.P.G. (2011). ss-TEA: Entropy based identification of receptor specific ligand binding residues from a multiple sequence alignment of class A GPCRs. BMC Bioinformatics 12 : 332.

Santangelo, E., Fonzo, V., Astolfi, S., Zuchi, S., Caccia, R., Mosconi, P., Mazzucato, A., and Soressi, G.P. (2003). The Cf-2 / Rcr3esc gene interaction in tomato (Lycopersicon esculentum) induces autonecrosis and triggers biochemical markers of oxidative burst at cellular level. Funct. Plant Biol. 30 : 1117.

Sarris, P.F., Cevik, V., Dagdas, G., Jones, J.D.G., and Krasileva, K.V. (2016). Comparative analysis of plant immune receptor architectures uncovers host proteins likely targeted by pathogens. BMC Biol. 14 : 8.

Saur, I.M., Bauer, S., Kracher, B., Lu, X., Franzeskakis, L., Müller, M.C., Sabelleck, B., Kümmel, F., Panstruga, R., Maekawa, T., and Schulze-Lefert, P. (2019). Multiple pairs of allelic MLA immune receptor-powdery mildew AVR effectors argue for a direct recognition mechanism. Elife 8.

Seong, K., Seo, E., Witek, K., Li, M., and Staskawicz, B. (2020). Evolution of NLR resistance genes with noncanonical N-terminal domains in wild tomato species. New Phytol.

Shao, Z.-Q., Xue, J.-Y., Wu, P., Zhang, Y.-M., Wu, Y., Hang, Y.-Y., Wang, B., and Chen, J.-Q. (2016). Large-Scale Analyses of Angiosperm Nucleotide-Binding Site-Leucine-Rich Repeat Genes Reveal Three Anciently Diverged Classes with Distinct Evolutionary Patterns. Plant Physiol. 170 : 2095–2109.

Shenkin, P.S., Erman, B., and Mastrandrea, L.D. (1991). Information-theoretical entropy as a measure of sequence variability. Proteins 11 : 297–313.

Stam, R., Nosenko, T., Hörger, A.C., Stephan, W., Seidel, M., Kuhn, J.M.M., Haberer, G., and Tellier, A. (2019a). The Reference Genome and Transcriptome Assemblies of the Wild Tomato Species Highlights Birth and Death of NLR Genes Between Tomato Species. G3 9 : 3933–3941.

Stam, R., Silva-Arias, G.A., and Tellier, A. (2019b). Subsets of NLR genes show differential signatures of adaptation during colonization of new habitats. New Phytol. 224 : 367–379.

Stewart, J.J., Lee, C.Y., Ibrahim, S., Watts, P., Shlomchik, M., Weigert, M., and Litwin, S. (1997). A Shannon entropy analysis of immunoglobulin and T cell receptor. Molecular Immunology 34 : 1067–1082.

Tamborski, J. and Krasileva, K.V. (2020). Evolution of Plant NLRs: From Natural History to Precise Modifications. Annu. Rev. Plant Biol. 71 : 355–378.

Van de Weyer, A.-L., Monteiro, F., Furzer, O.J., Nishimura, M.T., Cevik, V., Witek, K., Jones, J.D.G., Dangl, J.L., Weigel, D., and Bemm, F. (2019). A Species-Wide Inventory of NLR Genes and Alleles in Arabidopsis thaliana. Cell 178 : 1260–1272.e14.

Van Ghelder, C. and Esmenjaud, D. (2016). TNL genes in peach: insights into the post-LRR domain. BMC Genomics 17 : 317.

Wang, J., Hu, M., Wang, J., Qi, J., Han, Z., Wang, G., Qi, Y., Wang, H.-W., Zhou, J.-M., and Chai, J. (2019a). Reconstitution and structure of a plant NLR resistosome conferring immunity. Science 364.

Wang, J., Wang, J., Hu, M., Wu, S., Qi, J., Wang, G., Han, Z., Qi, Y., Gao, N., Wang, H.-W., Zhou, J.-M., and Chai, J. (2019b). Ligand-triggered allosteric ADP release primes a plant NLR complex. Science 364.

Wang, J., Zhang, L., Li, J., Lawton-Rauh, A., and Tian, D. (2011). Unusual signatures of highly adaptable R-loci in closely-related Arabidopsis species. Gene 482 : 24–33.

Wu, C.-H., Abd-El-Haliem, A., Bozkurt, T.O., Belhaj, K., Terauchi, R., Vossen, J.H., and Kamoun, S. (2017). NLR network mediates immunity to diverse plant pathogens. Proc. Natl. Acad. Sci. U. S. A. 114 : 8113–8118.

Yang, S., Li, J., Zhang, X., Zhang, Q., Huang, J., Chen, J.-Q., Hartl, D.L., and Tian, D. (2013). Rapidly evolving R genes in diverse grass species confer resistance to rice blast disease. Proc. Natl. Acad. Sci. U. S. A. 110 : 18572–18577.

Yang, Z. (2007). PAML 4: phylogenetic analysis by maximum likelihood. Mol. Biol. Evol. 24 : 1586–1591.

Zhang, P., Hiebert, C.W., McIntosh, R.A., McCallum, B.D., Thomas, J.B., Hoxha, S., Singh, D., and Bansal, U. (2016). The relationship of leaf rust resistance gene Lr13 and hybrid necrosis gene Ne2m on wheat chromosome 2BS. Theor. Appl. Genet. 129 : 485–493.

